# Generative design of functional disordered regions in a developmental transcription factor

**DOI:** 10.64898/2026.10.12.757977

**Authors:** Lance Denes, Ryan J. Emenecker, Nongyun Wang, Devi Chirumamilla, Rashmi Mohotti, Alex S. Holehouse, Liam Holt, Timothee Lionnet

## Abstract

Rapid advances in protein structure design have not translated to intrinsically disordered regions (IDRs). This is largely due to our incomplete understanding of how IDR sequence encodes function. Here, we use stem cell differentiation as a model to map functional sequence features in the C-terminal IDR of the transcription factor RUNX2, which is required to bind chromatin and regulate gene expression during osteogenesis. Surprisingly, linear IDR motifs are dispensable; instead, the conformational dynamics of the IDR predict function. We develop an algorithm for rational IDR design based on biophysical principles, and use it to generate functional IDRs *de novo*. These findings extend generative protein design beyond the folded proteome.

## Introduction

Advances in artificial intelligence (AI) have enabled prediction and design of folded proteins; however, these methods do not address the ∼35% of the human proteome that lack a well-defined structure (*1–3*). This is because the appropriate target for biologically functional IDR design is often ill-defined, as sequence-to-function models of IDRs have yet to be resolved and integrated (*4–9*).

IDRs encode biological function at different levels. First, short linear interaction motifs (SLiMs) (*10–12*), are precise sequences of ∼3-15 amino acids that can bind folded interactors via coordination of motif amino acids in structured binding pockets (*13*). However, recent work shows that chemically specific, multivalent interactions between IDRs and binding partner surfaces can augment and even replace motif-based interactions (*14*). These “fuzzy” complexes explain how transient, distributed IDR-structure or IDR-IDR interactions can give rise to well-defined molecular interactions (*15–20*). Molecular grammars – non-random amino acid compositions and patterning – have been proposed to define the chemically specific interactions that mediate such transient interactions, but how sequence grammar encodes interaction specificity remains an open question (*9*, *21–23*). Finally, the biophysical features of IDR conformational ensembles, for instance their global dimensions (e.g. the radius of gyration) and local dynamics (e.g. transient helicity or longer range intrachain contacts) also impact function (*3*, *24–28*). Each of these three regulatory layers imposes distinct constraints on IDR sequence space that can be distinguished by coupling rational sequence perturbation with functional assays (*7*).

Transcription factors (TFs) are particularly enriched for disordered regions; IDRs account for ∼55% of residues in human TFs (Fig. S1A) (*29*). Historically, TFs have been treated as modular units in which structured DNA-binding domains (DBDs) direct binding to specific DNA sequence motifs while flanking IDRs encode effector activity: the recruitment of co-factors that activate or repress nearby genes, or modify chromatin locally (*30*, *31*). This paradigm has guided large-scale, reductionist approaches to characterize either the DNA sequence motifs of isolated DBDs, or how diverse TF IDRs impact the expression of a reporter gene when recruited to its promoter by a constant DBD (*16*, *30*, *32–38*). While these approaches have been successful, mounting evidence suggests they are incomplete: First, TF IDRs can impact chromatin binding in cells: both IDR-IDR and IDR-DNA interactions have been proposed to guide DBDs to genomic binding sites (*39–42*). Second, effector activity screens that recruit IDRs to a reporter gene often do not generalize across promoters and cell types; some IDRs even flip between activation and repression depending on the context (*34*, *36*, *38*, *43*). Finally, while reporter gene assays have yielded models that predict general transcriptional activation domains found in IDRs from viral or stress TFs, we found that these models miss activity in many TFs, particularly those involved in development (Fig. S1B-C).

Developmental transcription factors typically regulate hundreds of genes, sometimes acting as both activators and repressors, all within a dynamic cellular context (*44*, *45*). These complex biological requirements mean that developmental TF IDR functions evolve under a large network of constraints unlikely to be entirely captured by studying a single reporter gene in a foreign cellular context. Indeed, the IDRs of several key TFs that direct skeletal development -including RUNX2, the master regulator of osteogenesis -were not active in any recent reporter screens despite previous reports of activating or repressing effector function in the literature (Fig. S1D-E) (*31*). We therefore sought to investigate the function of the RUNX2 IDR within its native context.

Here we developed a platform to measure integrated IDR function in a robust and quantitative differentiation phenotype, using osteoblast differentiation from mesenchymal stem cells driven by RUNX2 as a model. We show that the RUNX2 C-terminal IDR (CIDR) is required for differentiation and mediates RUNX2 association with chromatin. Despite near-perfect CIDR conservation to mice, we found that linear IDR motifs are dispensable for CIDR function. Using molecular dynamics simulations, we identified features of the CIDR conformational ensemble predicting its potential to drive differentiation. Finally, we used generative AI to design synthetic RUNX2 IDRs *de novo* with precise physicochemical features that complement WT activity despite almost no sequence identity.

Together, these findings demonstrate that specific biological function can be encoded in IDRs via rational design and provide a roadmap to probe IDR sequence space unexplored by nature.

## Results

### The RUNX2 CIDR is required for osteogenesis

RUNX2-regulated osteogenesis is an attractive model to study developmental TF IDR functions. First, RUNX2 loss-of-function leads to a dramatic phenotype: mice lacking RUNX2 fail to mineralize the skeleton, while in humans, RUNX2 mutations cause the congenital bone disease cleidocranial dysplasia (CCD) (*46–48*). Additionally, osteoblast differentiation from human mesenchymal stem cells (hMSC) is a well-characterized system that produces a robust and quantifiable mineralization phenotype *in vitro* (Fig. 1A) (*49*, *50*). Finally, RUNX2 contains a large, poorly characterized C-terminal IDR (CIDR), the truncation of which causes CCD (Fig. 1B) (*51*).

**Fig. 1.**
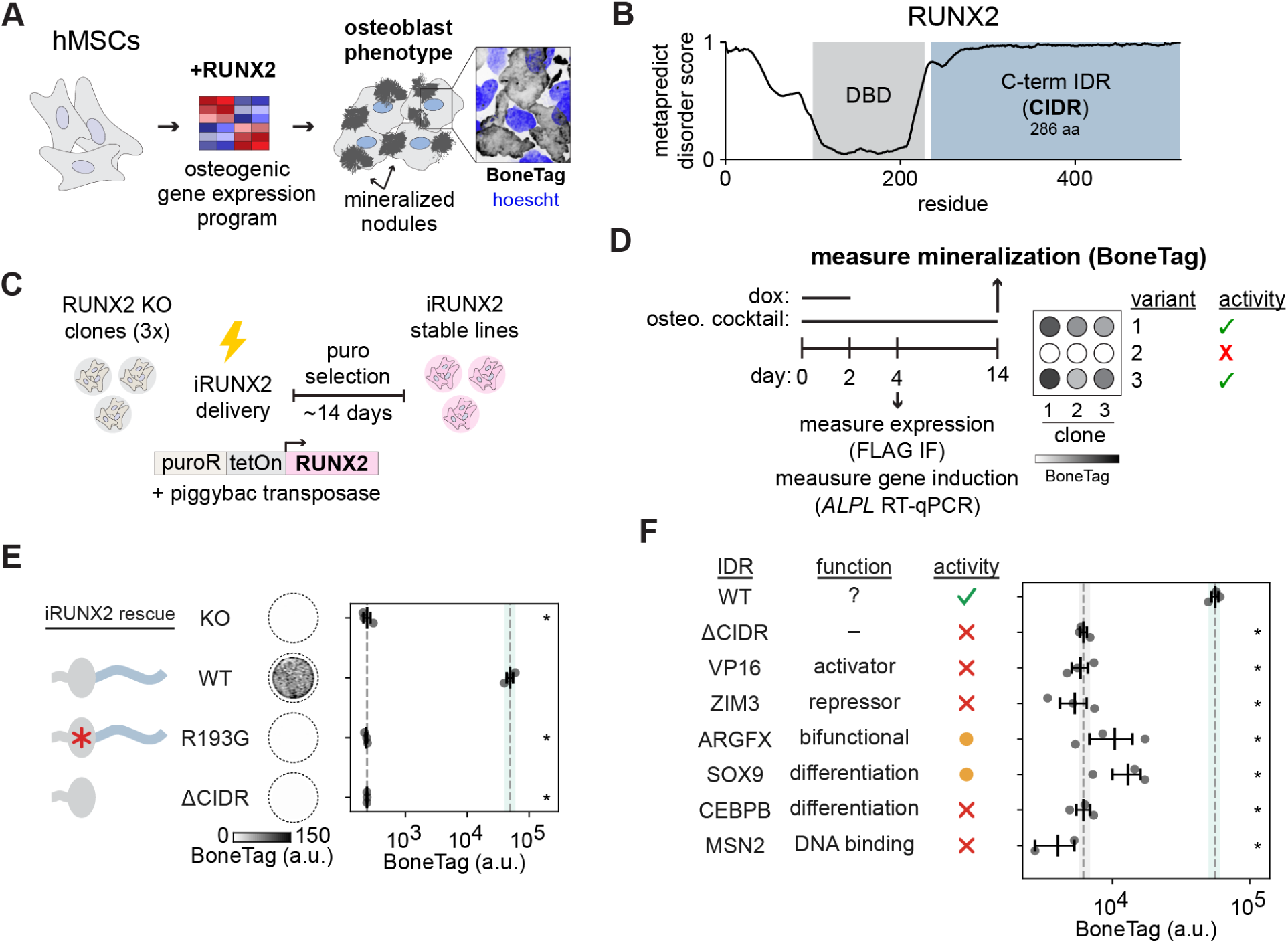
Osteoblast differentiation requires the RUNX2 CIDR. (**A**) Osteogenic differentiation of human mesenchymal stem cells (hMSCs) produces a quantifiable mineralization phenotype that depends on RUNX2 gene regulatory activity. (**B**) Metapredict disorder score across RUNX2. The structured DNA-binding domain (DBD, gray) is flanked by the C-terminal IDR (CIDR, blue). (**C**) Generation of inducible RUNX2 (iRUNX2) rescue lines. C Piggybac constructs are delivered to three independent RUNX2 knockout (KO) hMSCs clones by electroporation and selected under puromycin for ∼14 days to yield stable lines. (**D**) Triplicate wells are plated in parallel for qPCR, IF, and BoneTag mineralization assays. Cells are induced in osteoblast media with dox on day 0 and switched to osteoblast media without dox on from day 2 onwards. (**E**) Left: cartoon of iRUNX2 rescue constructs; center: representative BoneTag images; right: quantification of BoneTag signal. R193G is a DNA-binding domain loss-of-function mutation (red asterisk) from a cleidocranial dysplasia (CCD) patient. (**F**) We replaced the RUNX2 CIDR with IDRs drawn from various transcription factors and assayed osteogenesis as in E. From left to right: TF of origin of each IDR; annotated function;activity score; BoneTag quantification. In E and F, Each point represents the mineralization score from a single clone, averaged across three wells; error bars indicate SEM across clones. Dotted lines and shaded regions mark the mean and SD for WT and ΔCIDR. Statistics: one-way ANOVA on log10-transformed clone means followed by Dunnett’s multiple-comparison test against WT; *p < 0.05 vs. WT. No star indicates not significant vs. WT.

We developed an osteogenesis rescue assay to measure the activity of RUNX2 CIDR variants. We first generated RUNX2 knockout (KO) hMSC clones that could not differentiate into osteoblasts (Fig. S2A-E). We then generated RUNX2 rescue lines stably expressing inducible RUNX2 (iRUNX2) variants by piggybac integration of the transgenes into the RUNX2 KO clones (Fig. 1C). We induced RUNX2 expression with doxycycline and osteoblast differentiation with an established cocktail (see methods), and quantified the mineralization phenotype on day 14 by staining cells with 800CW BoneTag (a calcium-chelating compound conjugated to near-infrared fluorescent dye) and reading out well fluorescence (Fig. 1D). Importantly, we fused a FLAG tag to each of the RUNX2 variants to confirm consistent expression levels and nuclear localization on day 4 of differentiation via immunofluorescence, and measured induction of the osteoblast marker *ALPL* by RT-qPCR. We first benchmarked the system with WT human RUNX2, a nonfunctional DBD mutant identified in CCD patients (R193G) (*52*), and a deletion of the CIDR (ΔCIDR) (Fig. 1E). While WT iRUNX2 rescue cells produced a robust mineralization phenotype, neither R193G nor ΔCIDR rescues were distinguishable from KO, despite correct expression and localization of the variant (Fig. S3A-C), indicating the RUNX2 CIDR is required for osteoblast differentiation.

### Effector domain IDRs from other TFs do not complement RUNX2 CIDR function

We next tested whether RUNX2 variants containing a strong transcriptional activator (VP16) (*53*), a transcriptional repressor (ZIM3) (*34*), or a bifunctional domain (ARGFX, shown to activate or repress depending on the promoter) (*35*) in place of the CIDR could rescue osteogenic activity. Interestingly, although both the VP16 and ARGFX variants activated the osteoblast marker gene *ALPL* similar to WT RUNX2 early in differentiation, all three variants failed to rescue the mineralization phenotype (Fig. 1F, S4A-C). Only the ARGFX fusion showed measurable mineralization, but was inefficient compared to RUNX2 CIDR.

We reasoned that CIDR activity may depend on cellular context, i.e., sets of co-regulators or a chromatin environment specific to hMSCs. hMSCs are multipotent and can also differentiate into adipocytes or chondrocytes, driven by CEBPB and SOX9, respectively (*54*, *55*). Therefore, we reasoned that IDRs taken from these TFs may be functional in the cellular context of hMSCs.

However, IDRs from CEBPB and SOX9 also failed to fully rescue osteogenesis when fused to RUNX2 in place of the CIDR (Fig. 1F, S4A-C). These results suggest that the RUNX2 CIDR encodes distinct, osteogenic program-specific functional determinants.

Finally, previous work has implicated the yeast TF MSN2 IDR in dictating genomic binding (*40*, *56*). To investigate whether RUNX2 CIDR function could be complemented by an orthogonal chromatin-binding function, we tested it in our assay. However, substituting the RUNX2 CIDR for the MSN2 IDR did not rescue osteogenesis (Fig. 1F, S4A-C).

Together, these results demonstrate that orthogonal activation, repression, bifunctional or chromatin binding activities are insufficient to rescue the osteogenic activity of the RUNX2 CIDR. Thus the RUNX2 CIDR appears to encode a specific activity required for osteogenesis. We next sought to determine the sequence features underlying this activity.

### Degenerate local sequence features are sufficient for RUNX2 CIDR function

We suspected that the RUNX2 CIDR may encode a specific interaction in a short linear motif. In practice, these motifs mediate site-specific binding defined by intolerance to sequence scrambling, which would disrupt coordination of amino acids in a binding pocket, while avoiding the potential confounder of perturbing IDR composition (Fig. 2A) (*14*). To identify linear motifs within the RUNX2 CIDR (Fig. 2B), we divided the CIDR into 28 contiguous tiles of 8-11 amino acids and generated a series of RUNX2 variants in which each construct contained a scrambled version of one tile (Fig. S5A). To our surprise, all 28 variants successfully drove osteogenesis (Fig. 2C, S5B-C), including constructs whose scrambled tiles overlapped three previously characterized motifs with proposed molecular functions (*57*, *58*). These results indicate that CIDR function does not depend exclusively on a single linear motif.

**Fig. 2.**
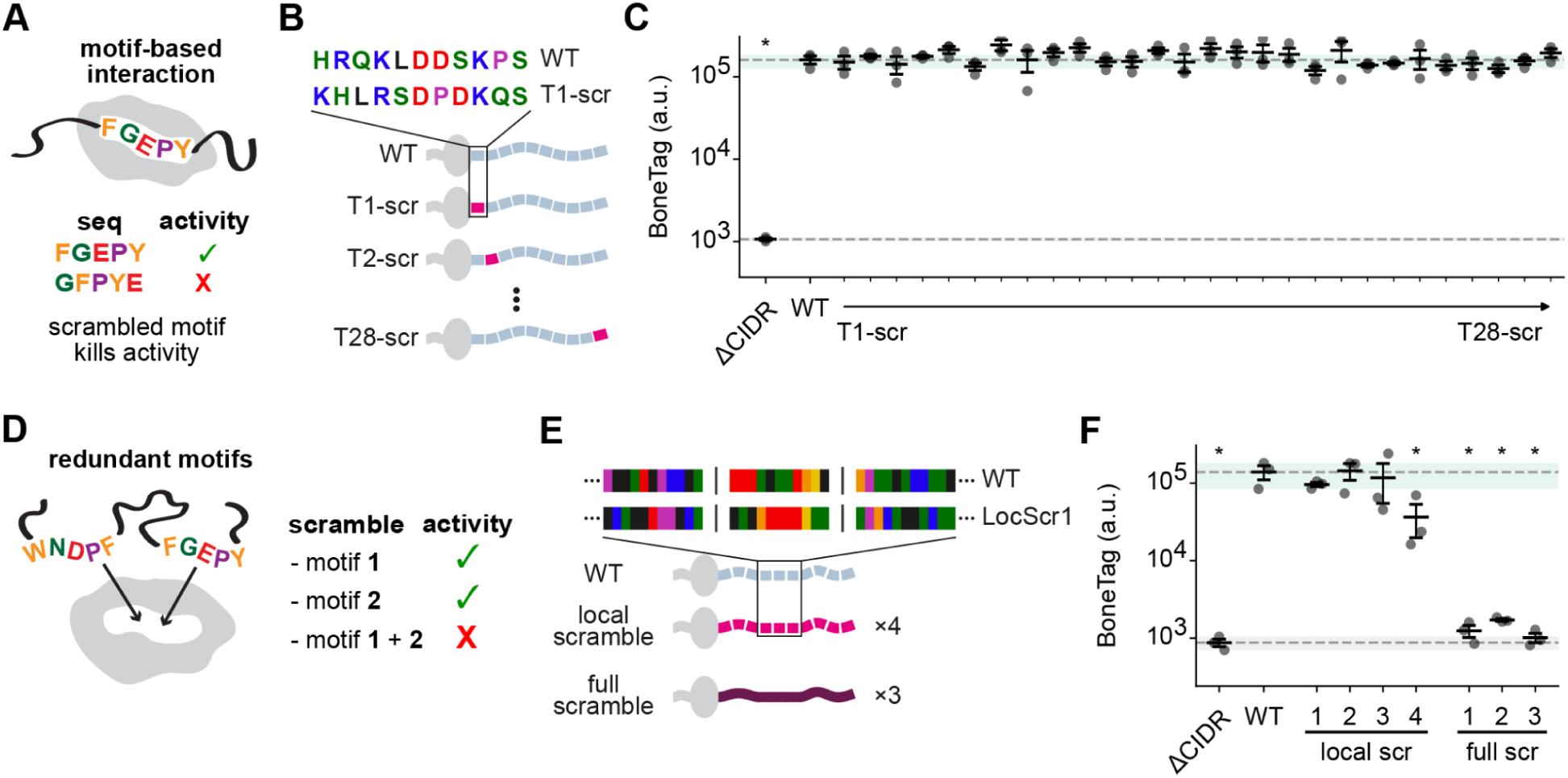
RUNX2 CIDR activity does not require sequence motifs but is sensitive to local amino acid composition. **(A)** Scramble mutations test whether linear interaction motifs are required for activity. (B) Design of tile scramble series. Each construct carries a scrambled version of a single 8-11 amino acid tile (magenta) in an otherwise wild-type (WT, gray) background. **(C)** Quantification of BoneTag mineralization assay for variants in (B). Each point represents the mineralization score from a single clone, averaged across three wells; error bars indicate SEM across clones. Dotted lines and shaded regions mark the mean and SD for WT and ΔCIDR. **(D)** Redundant motifs can compensate each others’ loss, requiring combined scrambling to reveal a phenotype. **(E)** Schematic of the local scramble strategy. Tiles from **(B)** were individually scrambled, then concatenated into a single “local scramble” (LocScr) construct. Four local scramble constructs were generated using different tile boundaries. Three full CIDR scrambles were generated by randomizing residue order across the entire CIDR, preserving its amino acid composition. **(F)** Quantification of BoneTag mineralization assay for the variants in (E). Points, error bars and reference lines as in (C). Statistics as in Fig. 1; *p < 0.05 vs. WT. No star indicates not significant vs. WT.

While individual tile scrambles rule out the essentiality of any one CIDR motif, there remains the possibility for motif redundancy, where loss of one motif is compensated by a different motif elsewhere in the IDR (Fig. 2D). Because testing all tile combinations was infeasible, we devised a strategy to test for motif redundancy (Fig. 2E). We concatenated all CIDR scrambled tiles into one sequence such that the entire sequence is scrambled, but only at a local level. Thus, the amino acids from every tile above are still constrained to remain within the 8-11 amino acid region that they came from, but they are no longer in the same order. We refer to these mutants as “local scrambles”. To our surprise, our first local scramble variant (LocScr1), (Fig. S6A) was functional (Fig. 2F, S7A). To confirm this result, we generated three additional local scramble CIDR variants by shifting the tile boundaries and independently scrambling the tiles. We also generated three “full scramble” variants in which we scrambled the entire CIDR sequence without restricting the positions of amino acids to their original windows (Fig. S6A). Each local scramble variant was functional, although one had a partial loss of function, while all full scramble variants were inactive (Fig. 2F, S7A-C).

Taken together, these results show that the RUNX2 CIDR encodes activity that – surprisingly – does not require its exact amino acid sequence order. However, CIDR requires some conservation of local amino acid chemical composition. These results imply that the physicochemical features of the CIDR play a crucial role in RUNX2 CIDR function.

### Both repulsive and attractive domains are required for RUNX2 CIDR function

We next investigated how the biophysical properties and conformational dynamics of the CIDR might relate to its function. IDRs are not fully disordered, but rather encode complex dynamical ensembles of partial structure. Recent advances in coarse-grained molecular dynamics simulations of IDRs (*59–62*) have revealed a rich, broad, sequence-encoded conformational landscape, enabling quantitative insights into how IDR sequences dictate intra-and inter-molecular interactions (*6*, *63–65*). We performed molecular dynamics simulations of the RUNX2 CIDR and quantified inter-residue contacts in the conformational ensemble (Fig. 3A, S8). We identified a “linker” / “patch” subdomain architecture in which four attractive patches (P1-4) were enriched for inter-residue contacts, while four repulsive linker regions (L1-4) were depleted of contacts (Fig. 3B). The patches were enriched for aromatic amino acids, which are known to drive inter-residue interactions in IDRs (*23*). Thus, molecular dynamics reveal subdomains with distinct physico-chemical properties that underlie the dynamic conformation of the CIDR.

**Fig. 3.**
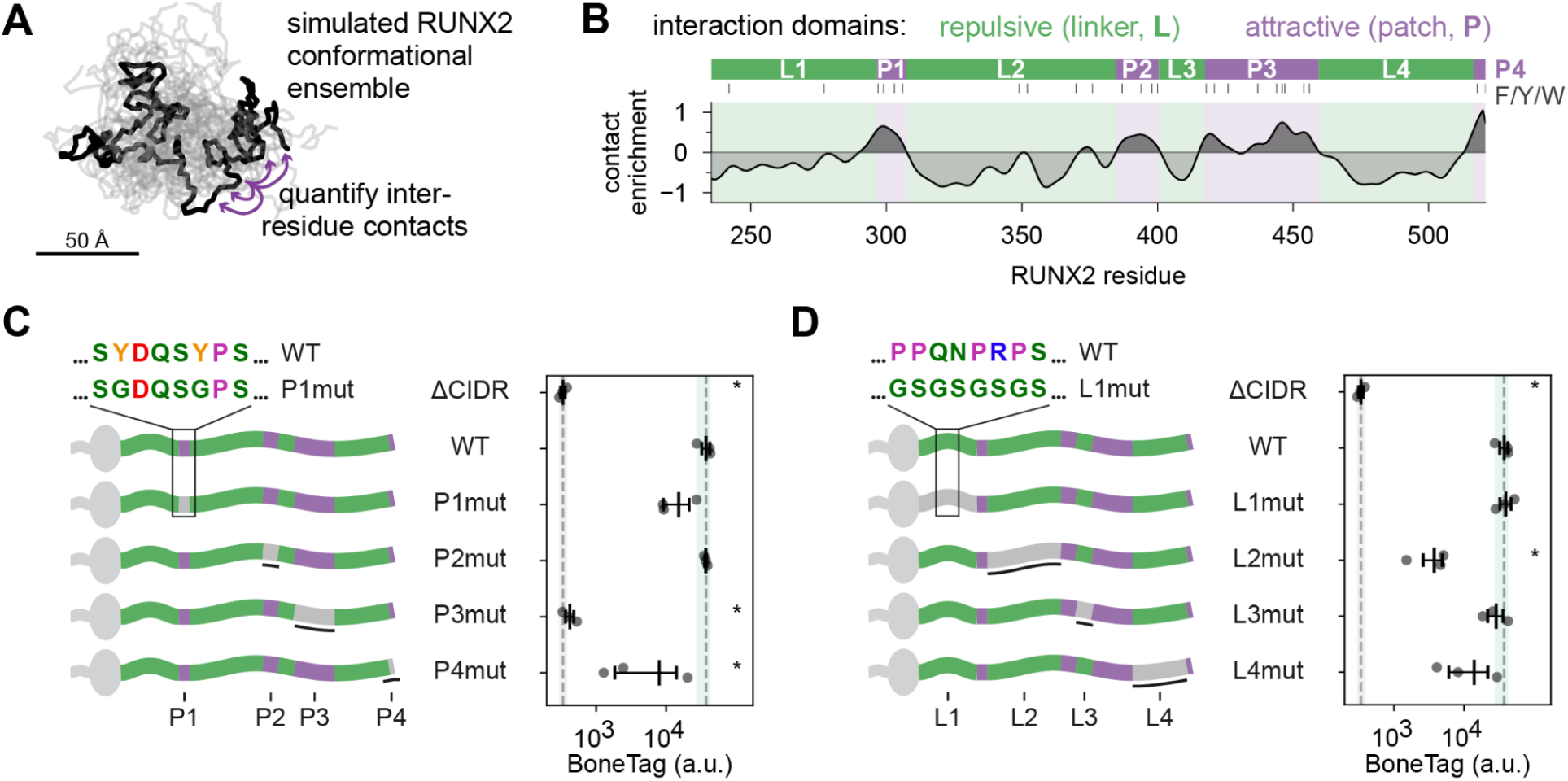
The RUNX2 CIDR is organized into alternating attractive and repulsive subdomains which are required for osteogenic activity. **(A)** Ten RUNX2 CIDR conformers (grey) with one conformer highlighted (black) from coarse-grained molecular dynamics simulation. **(B)** Inter-residue contact enrichment along the RUNX2 CIDR from simulations in (A). Contacts are normalized against the average inter-residue contact frequency as a function of sequence separation. Positive values mark four attractive patches (P1–P4, purple) and negative values mark four repulsive linkers (L1–L4, green). Tick marks indicate the positions of aromatic residues (F/Y/W), enriched within patches. **(C)** Schematic of attractive patch mutants and quantification of corresponding BoneTag mineralization assay. Aromatic residues in each patch were mutated to glycines. **(D)** Schematic of repulsive linker mutants and quantification of corresponding BoneTag mineralization assay. Linker residues in each patch were mutated to glycine-serine repeats of matched length. In C and D, Each point represents the mineralization score from a single clone, averaged across three wells; error bars indicate SEM across clones. Dotted lines and shaded regions mark the mean and SD for WT and ΔCIDR. Statistics as in Fig. 1; *p < 0.05 vs. WT. No star indicates not significant vs. WT.

To test the requirement of linker and patch subdomains for RUNX2 CIDR function, we generated CIDR variants that neutralized the repulsive interactions in each linker with polar Glycine-Serine repeats of the same length (L1mut, L2mut, L3mut, L4mut), and variants that neutralized the attractive interactions in each patch by replacing aromatic residues with glycines (P1mut, P2mut, P3mut, P4mut). Interestingly, these variants showed a range of osteogenic potential. Among patch mutants, P1mut and P2mut were able to support wild-type levels of mineralization, P4mut showed a hypomorphic phenotype, and P3mut was completely nonfunctional (Fig. 3C, S9). While L1mut and L3mut fully rescued mineralization, L2mut and L4mut RUNX2 showed a strong hypomorphic phenotype (Fig. 3D, S9).

Together, these results show that the RUNX2 CIDR contains both attractive and repulsive subdomains that are required for its function, possibly explaining why its sequence cannot be completely reordered, but local chemical properties must be maintained. As L2mut and P3mut led to severe differentiation disruption despite opposing effects on the conformational ensemble (neutralization of respectively a repulsive vs. an attractive subdomain), we chose to further characterize the molecular mechanisms of loss of CIDR function in these two mutants.

### RUNX2 CIDR is required to bind motif clusters in inaccessible chromatin

A preponderance of literature focuses on the hypothesis that TF IDRs recruit co-regulators to the DNA-bound TF. However, several recent studies have shown that TF IDRs can also impact chromatin binding in cells (Fig. 4A) (*39*, *40*, *42*). We thus hypothesized that the RUNX2 CIDR might impact chromatin binding.

**Fig. 4.**
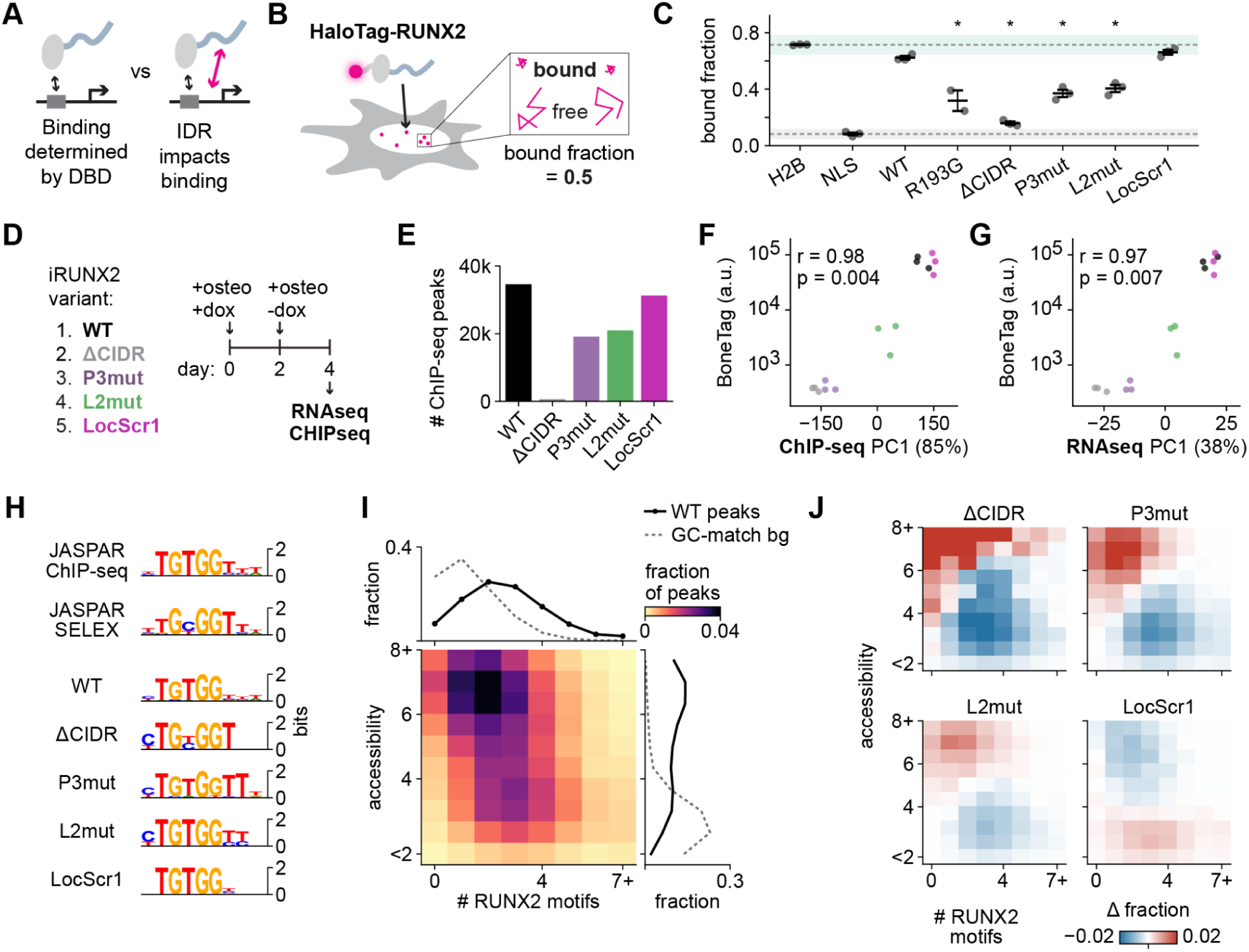
The RUNX2 CIDR directs DNA binding to motif clusters in inaccessible chromatin.(A) Two models for CIDR function: binding by the DNA-binding domain alone (left) or CIDR-modulated (right). **(B)** Single-molecule tracking in hMSC lines expressing RUNX2-HaloTag fusions; molecules classified as chromatin-bound or freely diffusing by mobility (Methods). **(C)** Chromatin-bound fraction for RUNX2 variants and controls. Points, per-experiment cell averages; error bars, SD across three experiments. Dotted lines and shading, H2B and NLS control mean and SD. Statistics as in Fig. 1; *p < 0.05 vs. WT. **(D)** ChIP-seq and RNA-seq experimental design in iRUNX2 rescue lines. **(E)** Number of ChIP-seq peaks per variant. **(F and G)** Mineralization activity (BoneTag) against PC1 of differentially bound peaks **(F)** or differentially expressed genes **(G)** relative to WT; Pearson *r* on construct means. **(H)** Top *de novo* motif in peaks bound by each variant; JASPAR RUNX2 SELEX and ChIP-seq motifs for reference (*37*, *68*). (I) Joint distribution of chromatin accessibility, DNase-seq FPKM (*49*), and RUNX2 motif count (FIMO) within WT peaks; 1D histograms above and right; dotted line, GC-matched background. (J) Percent change in peak fraction per accessibility × motif count bin, variant versus WT.

We first tested this hypothesis by direct imaging of RUNX2. Single molecule tracking (SMT) directly measures the binding of TFs to chromatin in living cells by measuring the fraction of freely diffusing to near-stationary molecules in the nucleus (Fig. 4B S10); it has the advantage over genomics approaches of capturing overall chromatin-TF association, i.e. both transient chromatin-scanning episodes and motif-binding events are captured (*66*). We generated hMSC lines expressing HaloTag fusions of RUNX2 WT, R193G (loss of function CCD mutation in the DBD), ΔCIDR, LocScr1, L2mut, and P3mut and performed SMT (Fig S10A-C). We also generated control lines expressing H2B-and NLS-HaloTag fusions as controls for chromatin-bound and free molecules. For both functional RUNX2 variants, WT and LocScr1, the slow-diffusing fraction was near the H2B control, indicating most molecules were chromatin-bound (Fig. 4C). Surprisingly, deleting the CIDR effectively abolished binding; this effect was even stronger than the R193G DBD mutant. L2mut and P3mut exhibited chromatin occupancy intermediate between those of RUNX2 WT and ΔCIDR. These results indicate that the RUNX2 CIDR is required for chromatin binding, and that perturbations to either repulsive linker or attractive patch subdomains impair this behavior.

We next performed RNAseq and ChIP-seq on differentiating cells expressing the same set of RUNX2 CIDR variants to determine whether genome-wide chromatin binding and gene regulation were impacted (Fig. 4D). Consistent with SMT data, the ΔCIDR variant lost the majority of ChIP-seq peaks, L2mut and P3mut showed a partial loss, and LocScr1 bound a similar number of sites relative to WT (Fig. 4E). The first component (PC1) from a principal components analysis (PCA) on differentially bound peaks was strongly correlated with mineralization phenotype (Fig. 4F, Fig. S11A). This was also true of the PCA on differentially expressed genes in RNAseq analyses (Fig. 4G, S11B), which were enriched for genes involved in osteogenesis (Fig S11C-E), supporting a direct correlation between loss of chromatin association and failure to differentiate. *De novo* DNA motif discovery revealed that each variant bound the consensus RUNX2 motif, though with slight variations (Fig. 4H) (*67*). The RUNX2 DNA motif derived from *in vitro* SELEX experiments on the isolated DBD differs from the

ChIP-seq motif in the underspecification of a central T (*37*, *68*). The ΔCIDR motif matched the SELEX-derived motif while other variants matched the ChIP-seq motif, suggesting that this slight motif variation is caused by the presence of CIDR rather than experimental context.

Flanking residues were also more strongly specified in the binding-deficient CIDR variants, suggesting that the CIDR can facilitate access to weaker motifs.

As all CIDR variants bound the consensus RUNX2 motif, we asked whether other genomic features were associated with differential binding. We found mild enrichment for CEBPB and KLF5 motifs in peaks gained by ΔCIDR, P3mut, and L2mut variants (Fig. S12A). However, the strongest signal in peaks lost in variants was for RUNX2 motifs. This indicated that binding defects were unlikely to result from a lost interaction with a preferred sequence-specific TF partner. To determine whether chromatin state impacted CIDR-dependent binding, we analyzed DNase accessibility in undifferentiated hMSCs prior to the onset of RUNX2 activity (*49*). When we compiled DNase accessibility at WT RUNX2 ChIP-seq peaks along with the RUNX2 motif count under those peaks, we noticed two populations: high accessibility, low motif count *vs.* low accessibility, high motif count (Fig. 4I). ΔCIDR, L2mut, and P3mut lost binding at low accessibility, high motif count sites and gained binding at high accessibility, low motif sites (Fig. 4J, Fig. S13A). LocScr1, on the other hand, bound more to low accessibility sites irrespective of motif count. A linear model showed motif count and accessibility were both predictive of differential ΔCIDR, L2mut, and P3mut binding, whereas only accessibility was predictive of differential LocScr1 binding (Fig. S13B). Together, our data suggest a model in which the RUNX2 CIDR is required for RUNX2 to bind sites with clustered motifs in less accessible chromatin to drive the osteogenic gene expression program.

### *De novo* design of functional RUNX2 CIDRs

Because our data demonstrate that linear IDR motifs are not required for RUNX2 CIDR activity, we wondered whether we could instead leverage sequence-encoded biophysical properties to design functional synthetic IDRs *de novo*. We compared the conformational ensembles obtained from molecular dynamics simulations on the scrambled, linker, and patch mutants to the WT CIDR ensemble. Divergence from the WT ensemble scaled inversely with mineralization activity (Fig. 5A-B, S14A-C). This suggested that the RUNX2 CIDR conformational dynamics could provide a tractable design target to produce functional sequences.

**Fig. 5.**
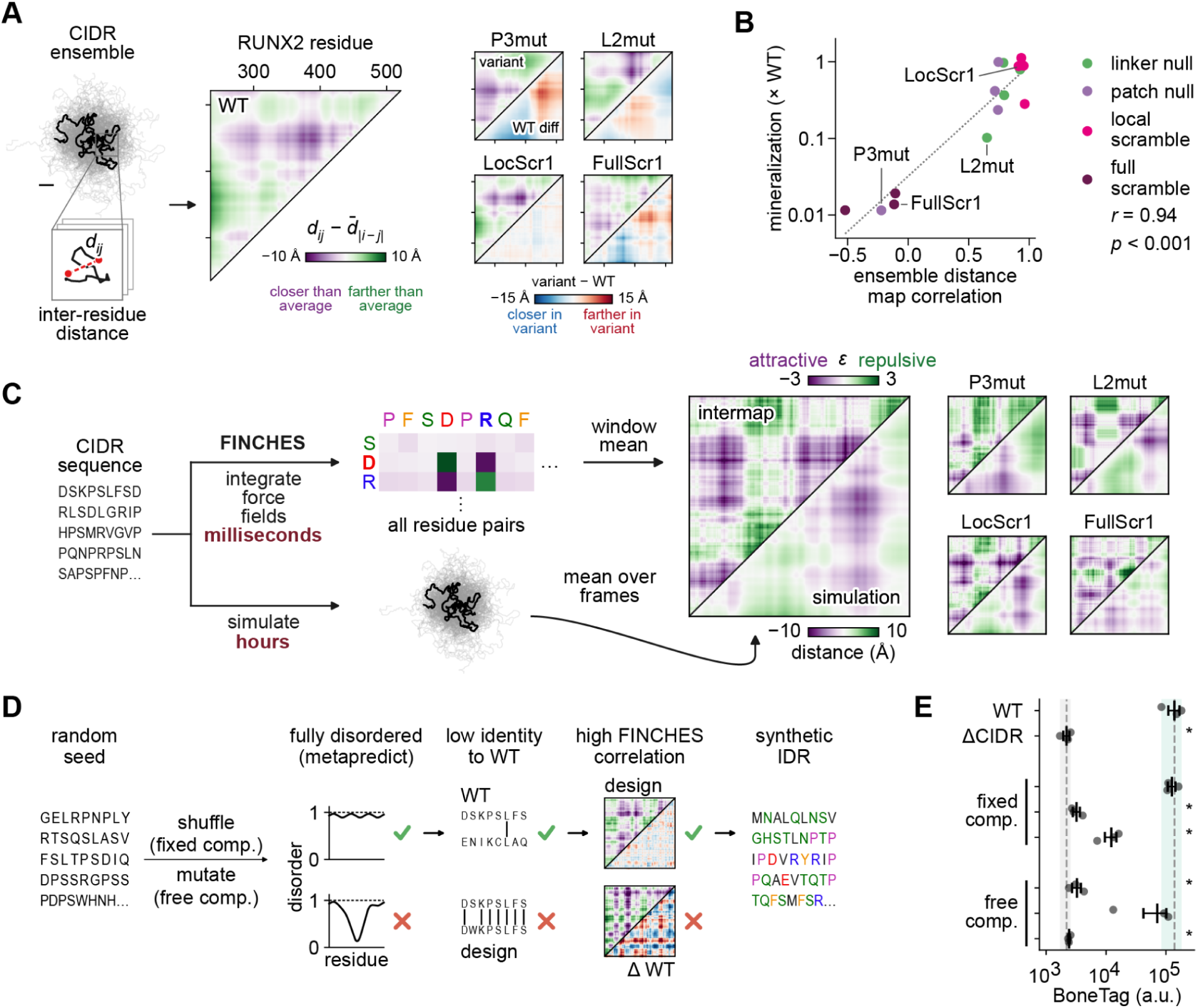
Generative design of functional RUNX2 CIDRs. **(A)** Left, schematic of the interresidue distance calculation from simulations of the WT RUNX2 CIDR; mean interresidue distances across frames are normalized to the mean at matched sequence separation. Right, distance maps for each variant and their difference from WT CIDR. **(B)** Batch-normalized mineralization activity versus the correlation of each variant distance map with WT CIDR. **(C)** FINCHES intermap strategy, which estimates mean field interaction strengths between sequence windows by integrating amino acid force fields over space. Right, distance maps from simulations compared with FINCHES intermaps for each variant. **(D)** Generative IDR design algorithm.

IDR conformational ensembles are driven by non-covalent chemical interactions between amino acid side chains (*3*, *69–73*). Our recently published tool FINCHES uses the non-bonded interaction strength parameters from molecular dynamics force fields to estimate chemical attractiveness between subdomains of arbitrary IDR sequences (*8*). This is a highly efficient method to construct an interaction map (intermap) for an IDR sequence as a proxy for the ensemble dynamics. This computational efficiency, generating intermaps in milliseconds, enables us to use intermap similarity as a generative design target to approximate constraints on the conformational ensemble (Fig. 5C).

To design *de novo* IDRs with CIDR-like ensembles, we combined FINCHES with our IDR design tool, GOOSE (*7*). Using FINCHES intermaps as a design constraint within GOOSE’s SequenceOptimizer functionality, we can design synthetic IDRs to match a desired FINCHES intermap along with any number of additional constraints. Specifically, we generated *de novo* IDRs satisfying the following constraints: maximal disorder, minimal sequence identity to WT CIDR (to avoid the trivial result of recapturing natural sequences), and maximal correlation to the WT CIDR FINCHES intermap (Fig. 5D). We generated one batch of sequences with fixed amino acid composition (i.e. same amino acid proportions as CIDR) and one with free composition (i.e. arbitrary amino acids). We expressed six variants (three in each category) and confirmed they localized to the nucleus similarly to WT RUNX2. Remarkably, two synthetic variants -one of fixed and one of free composition -were fully functional, matching the success rates reported for state-of-the-art structured binder design (Fig. 5E) (*74*). These results establish that *de novo* IDRs built using the conformational ensemble as a design blueprint can perform the work of a native effector domain and drive cell differentiation.

## Discussion

Here, we used stem cell differentiation as a model to map the functional sequence elements of a TF IDR. We showed that features of the RUNX2 CIDR conformational ensemble predict osteogenic activity, while linear sequence motifs are dispensable. We identified ensemble CIDR features required for binding to chromatin and generated synthetic IDR variants *de novo* using a generative design algorithm built on these biophysical principles.

While small linear motifs mediate IDR function in many contexts, reductionist assays have suggested that degenerate sequence features (enrichment for acidity and hydrophobicity) are sufficient for TF IDRs to activate a reporter gene (*16*, *20*, *75–77*). However, we found that these classical TF activation domains are insufficient for the more precise function of driving in vitro osteoblast differentiation, which requires the RUNX2 CIDR. Nevertheless, direct test with shuffle variants also revealed that site-specific binding motifs are dispensable for RUNX2 CIDR function, assayed here in its native molecular and cellular context. Our work thus establishes that despite 99% human-mouse CIDR conservation, *de novo* CIDR sequences designed to match the WT conformational ensemble are competent to orchestrate an entire differentiation program. We nevertheless note that, given we only probed *in vitro* differentiation and given RUNX2 has pleiotropic functions (*78–80*), we cannot exclude the possibility that the CIDR function might require linear motifs in other contexts. Moreover, we suggest that this potential ability to multiplex chemically specific (as reported here) and site-specific (motif-based) interactions may be key to enabling IDRs to engage in multiple orthogonal modes of molecular and cellular function.

Interestingly, all locally scrambled CIDR variants tested here were active, while several *de novo* designs were completely dead. This suggests that functional CIDR sequence features are captured by local amino acid composition, but not by the conformational ensemble alone. One likely explanation for this is that the coarse-grained models imperfectly capture IDR interactions and conformational biases. Another non-mutually exclusive possibility is that redundant, degenerate motifs are preserved at some frequency by local scramble designs, while they are unlikely to emerge in a synthetic sequence by chance. Finally, the conformational ensemble might confer a permissive 3D context for fuzzy interactions which rely on specific residues, thus composition-specific. This would explain why local scramble variants -which conserve both conformational ensemble and local composition -preserve function. On the other hand, some synthetic variants, while retaining the correct conformational ensemble, might lack key residues in their required context, and thus be defective. Distinguishing between these possibilities will require better biophysical models and high-throughput functional screens capable of identifying degenerate linear sequence features. Nevertheless, our findings suggest that IDR conformational dynamics are a key gatekeeper of function, and that taking this role into account will improve synthetic IDR design. For example, if a new synthetic motif inadvertently disrupts conformational dynamics, this could lead to an unexpected loss of function. Our approach can help avoid these pitfalls, i.e. by allowing the insertion of a motif of interest to the position where it minimally disrupts the conformational ensemble of the surrounding IDR.

Our work adds to a growing body of evidence that IDRs are required for many TFs to effectively bind chromatin in cells (*41*, *81*). While the ΔCIDR RUNX2 variant could still bind its motif, it was only found in the most open chromatin regions. How the CIDR potentiates binding to less accessible chromatin is an open question; mechanisms ranging from direct DNA contacts to interactions with protein partners have been proposed for other TFs (*41*). Since CIDR favors RUNX2 association with multiple RUNX2 motifs, it is possible that the CIDR might mediate homotypic interactions between RUNX2 molecules, thus increasing RUNX2 avidity at motif clusters (*82*). Regulatory specificity may thus emerge from two matched codes: the physicochemical patterning of the CIDR (which sets what interactions are possible) and the motif architecture of the genome (which sets where those interactions can occur).

Decades of structural biology are built on the axiom that sequence encodes structure, and structure equals function (*83*). This vision has culminated in AI-accelerated protein structure design, making the *de novo* design of protein binders routine and possible even in non-expert labs. Yet, since the mechanisms relating sequence to function are still being defined in the unstructured proteome (*3*), IDR design has lagged behind (*84*). Our ensemble-dynamics guided framework extends *de novo* protein engineering into the disordered proteome, yielding synthetic IDRs that support cell differentiation, and open the door to encoding new functions for stem cell reprogramming.

Fixed-composition designs iteratively scramble the RUNX2 CIDR sequence, optimizing for full disorder, low sequence identity to WT, and high FINCHES intermap correlation.

Free-composition designs mutate individual residues each iteration. **(E)** BoneTag mineralization assay for de novo IDRs. Dots are individual clones, each averaged across three wells; error bars are the SEM across clones. Dotted lines and shaded regions are the mean and SD for WT and ΔCIDR. Statistics as in Fig. 1; *p < 0.05 vs. WT. No star indicates not significant vs. WT.

## Acknowledgments

We thank current as well as past members of the Lionnet, Holt, and Holehouse labs as well as other labs in the Institute for System Genetics at the NYU School of Medicine for feedback and discussions while the work was in progress. We thank Esteban Mazzoni and Omar Qassab for help with CHIPseq methodology, Peter Whitney, Nestor Saiz, and Jacob Swanson for advice on the manuscript, Grace Avecilla for discussions on RUNX2 evolutionary genetics, and Shahar Sukenik for discussions on IDR ensemble function. We also thank NYU Grossman School of Medicine’s Genome Technology Center (RRID: SCR_017929) for sequencing services and the High-Performance Computing Facility of the Center for Health Informatics and Bioinformatics at the NYU Grossman School of Medicine for compute resources.

## Author contributions

Conceptualization: LD, LH, TL; Formal analysis: LD, RE, AH; Funding acquisition: AH, LH, TL; Investigation: LD, NW, DC, RM; Methodology: LD, RE; Supervision: AH, LH, TL; Visualization: LD, RE, AH, LH, TL; Writing – original draft: LD; Writing – review & editing: RE, AH, LH, TL.

## Funding

TL is supported by NIH grants R01GM149835 and R01AG075272. ASH and RJE are supported by the National Cancer Institute at the National Institutes of Health via DP2-CA290639. NYU Langone’s Genome Technology Center is partially supported by Cancer Center Support Grant P30CA016087 at the Laura and Isaac Perlmutter Cancer Center.

## Competing interests

TL holds intellectual property rights related to Janelia Fluor dyes used in this publication.

## Data, code, and materials availability

Sequencing data have been submitted to the Gene Expression Omnibus (GEO). Code used for molecular dynamics simulations, single molecule tracking, genomics, and IDR generative design is available on Github (timotheelionnet/RUNX2_generative_design). Single molecule tracking and molecular dynamics data are available through Zenodo. Cell lines generated are available upon request.

## Supplementary material

**Fig. S1.**
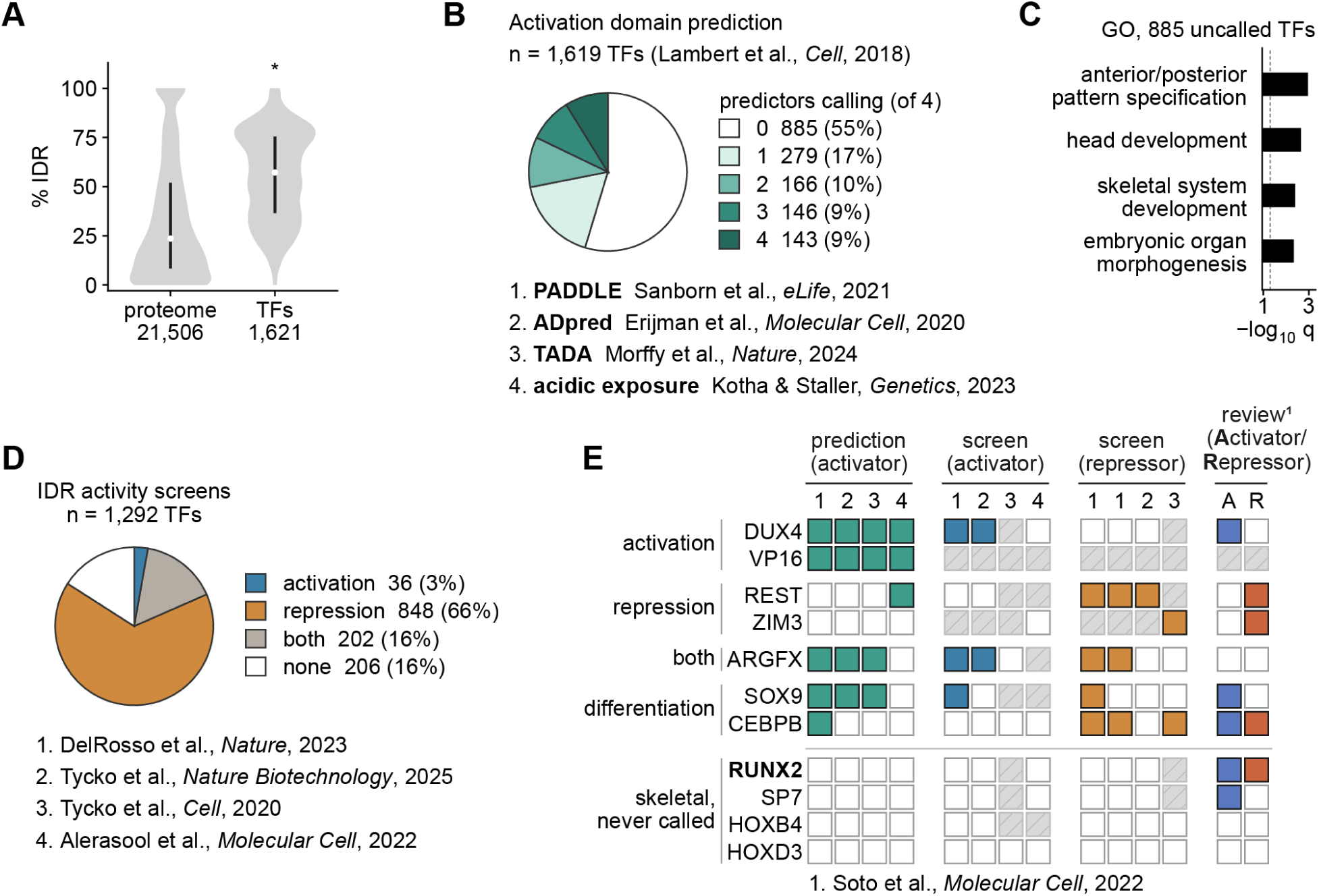
Regulatory activity in vertebrate development TFs is missed by reporter gene approaches. **(A)** Percent of residues in IDRs in the human proteome (n = 21,506 reviewed proteins, which includes the TFs) vs. the human TFome (n = 1,621 Lambert TFs). Disorder predicted using metapredict. Violins are densities scaled to their own maximum. Black bars are the interquartile range and white dots are the median. *p < 0.05, IDR fraction in TFs vs. non-TF, Mann–Whitney U test. **(B)** Fraction of TFome predicted to contain activation domains by each of four independent model families (PADDLE, ADpred, TADA and acidic exposure), one call per family; the exact rule used for each is given in Materials and methods. **(C)** Gene Ontology terms enriched among the 885 TFs called by no predictor, as −log10 q (Fisher’s exact test, Benjamini–Hochberg correction, background = the 1,619 scored TFs). Dashed line is q = 0.05. **(D)** Fraction of TFs (n = 1,292) called as activators, repressors, both, or neither across four IDR activity screens. **(E)** Per-TF calls for example TFs across activation-domain predictors, activator and repressor screens, and a literature review. Filled cells are calls, empty cells were tested and not called, and hatched cells were not tested. RUNX2 and other skeletal TFs are never called.

**Fig. S2.**
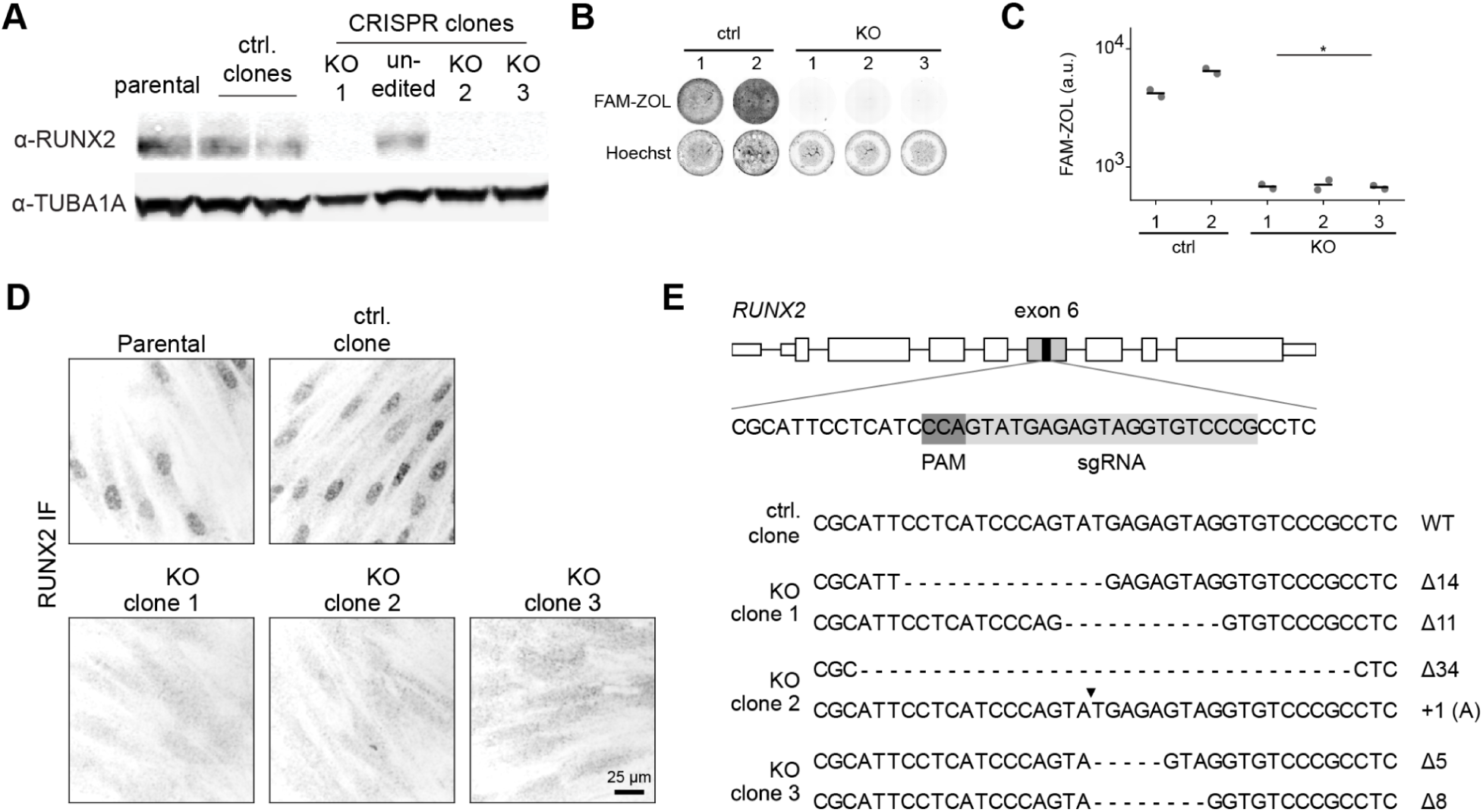
RUNX2 KO MSC generation. **(A)** Immunoblot for RUNX2 and TUBA1A (loading control) in parental MSCs, control clones, and clones that received CRISPR/Cas9 RNPs targeting RUNX2. **(B)** Whole-well FAM-ZOL mineralization and Hoechst images of control and RUNX2 KO clones on day 14 of differentiation. **(C)** Quantification of FAM-ZOL signal in (B). Dots are individual wells (two wells per clone); lines are the mean of wells. **(D)** RUNX2 immunofluorescence on day 4 of differentiation in parental MSCs, a control MSC clone, and three RUNX2 KO clones. Scale bar, 25 µm. **(E)** Schematic of gRNA targeting exon 6 of RUNX2 and sequenced alleles of each clone. In (C), *p = 0.002, KO vs. control, unpaired two-tailed t-test on log10-transformed clone means; n = 2 control and 3 KO clones.

**Fig. S3.**
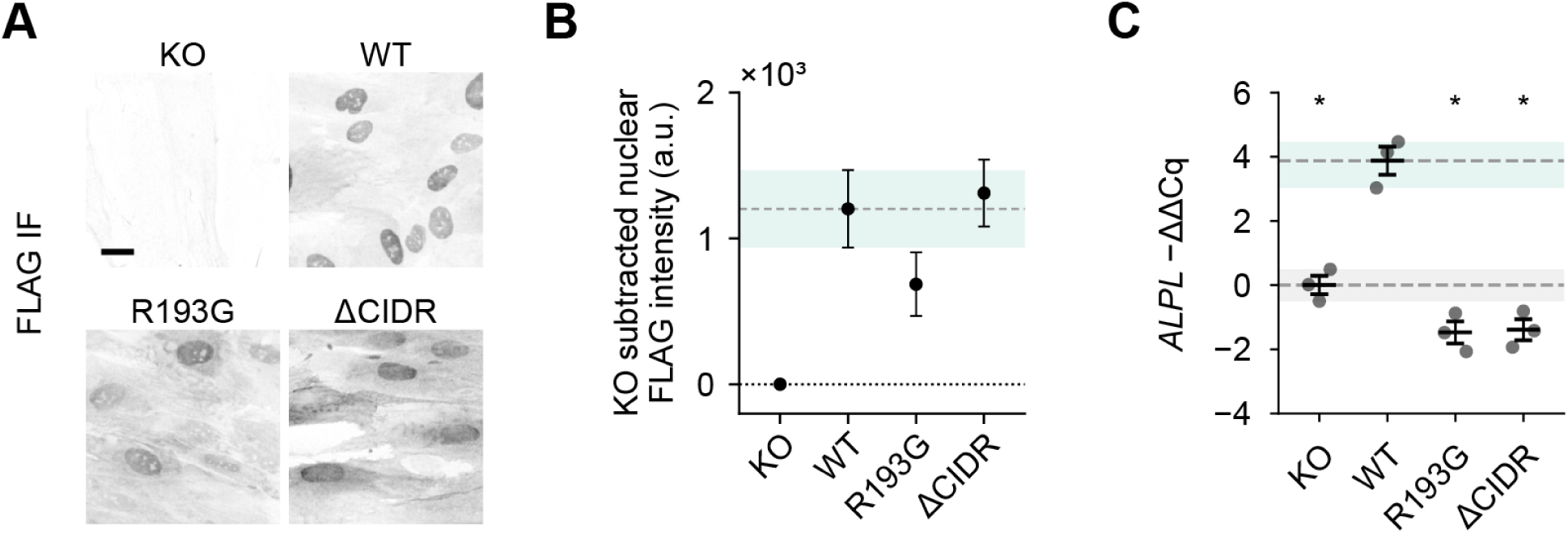
Expression and *ALPL* induction in RUNX2 control and CIDR deletion variants. **(A)** Anti-FLAG immunofluorescence on day 4 of osteoblast differentiation in KO, WT, R193G and ΔCIDR iRUNX2 rescue lines. Scale bar 25 µm. **(B)** KO-subtracted nuclear intensity of cells in (A). Dots and error bars are the mean and s.d. of each clone’s mean nuclear intensity. Shaded region and dashed line are WT mean and s.d.; dotted line is the KO floor. **(C)** *ALPL* induction by RT-qPCR on day 4 of osteoblast differentiation, as −ΔΔCq relative to KO (ΔCq = Cq*_ALPL_* − Cq*_TBP_*; higher values indicate more *ALPL*). R193G was measured on a separate plate and normalised through the ΔCIDR clones shared by both plates. *TBP* was used as the reference gene. Grey dots are clones (n = 3 per variant). Error bars are mean and SEM of clones. Green and grey shaded regions are the WT and KO clone ranges, with dashed lines at their means. Statistics: one-way ANOVA on log10-transformed clone means followed by Dunnett’s multiple-comparison test against WT; *p < 0.05 vs. WT. No star indicates not significant vs. WT.

**Fig. S4.**
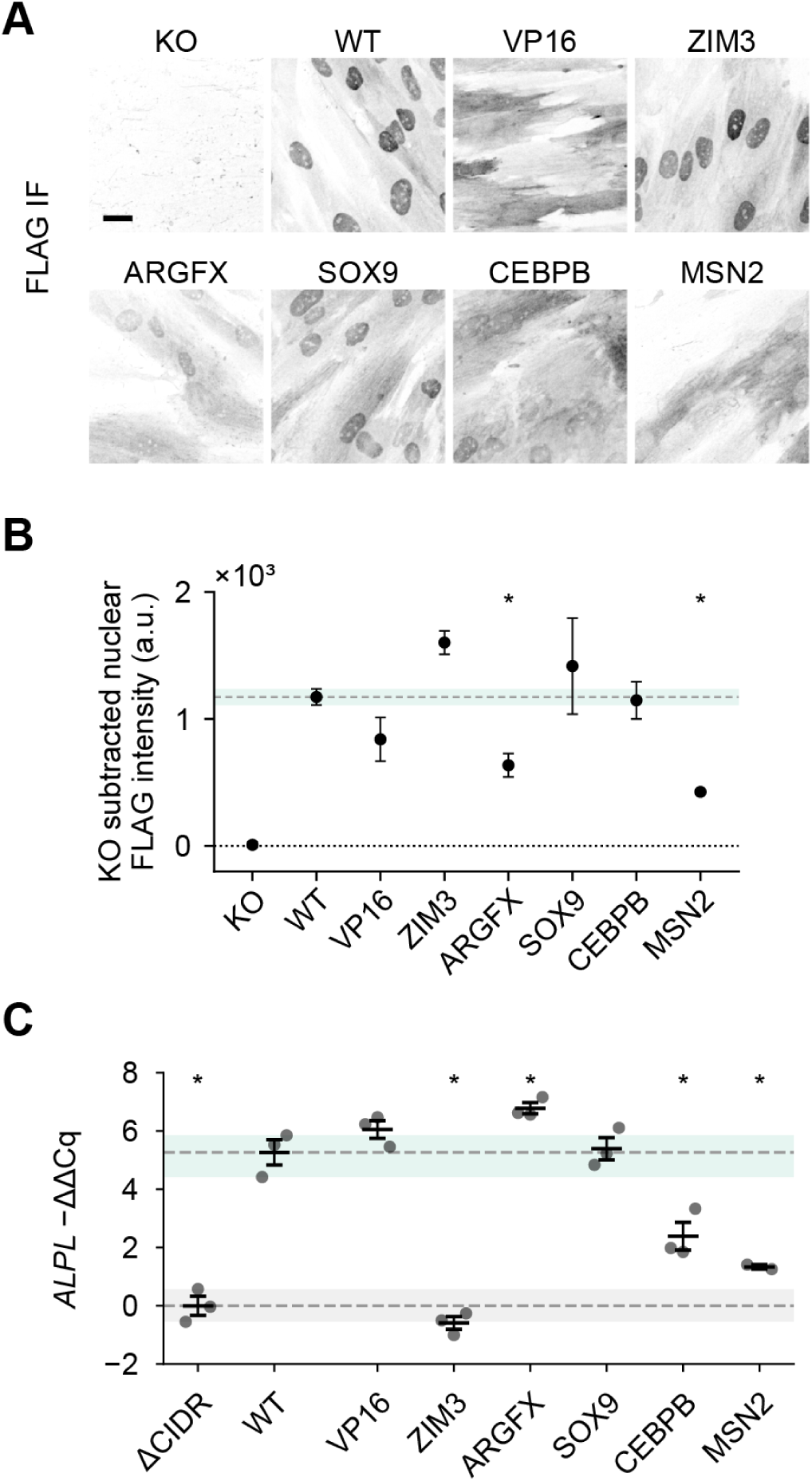
Expression and *ALPL* induction in RUNX2 IDR chimeras. **(A)** Anti-FLAG immunofluorescence on day 4 of osteoblast differentiation in KO, WT, and VP16, ZIM3, ARGFX, SOX9, CEBPB and MSN2 IDR chimera iRUNX2 rescue lines. Scale bar 20 µm. **(B)** KO-subtracted nuclear intensity of cells in (A), quantified as in Fig. S3B. Dots and error bars are the mean and s.d. of each clone’s mean nuclear intensity (n = 2 clones for MSN2). Shaded region and dashed line are WT mean and s.d.; dotted line is the KO floor. **(C)** *ALPL* induction by RT-qPCR on day 4 of osteoblast differentiation, as −ΔΔCq relative to ΔCIDR (ΔCq = Cq*_ALPL_* − Cq*_TBP_*; higher values indicate more *ALPL*). *TBP* was used as the reference gene. Grey dots are clones (n = 3 per variant; n = 2 for MSN2). Error bars are mean and SEM of clones. Green and grey shaded regions are the WT and ΔCIDR clone ranges, with dashed lines at their means. Statistics as in Fig. S3. *p < 0.05 vs. WT. No star indicates not significant vs. WT.

**Fig. S5.**
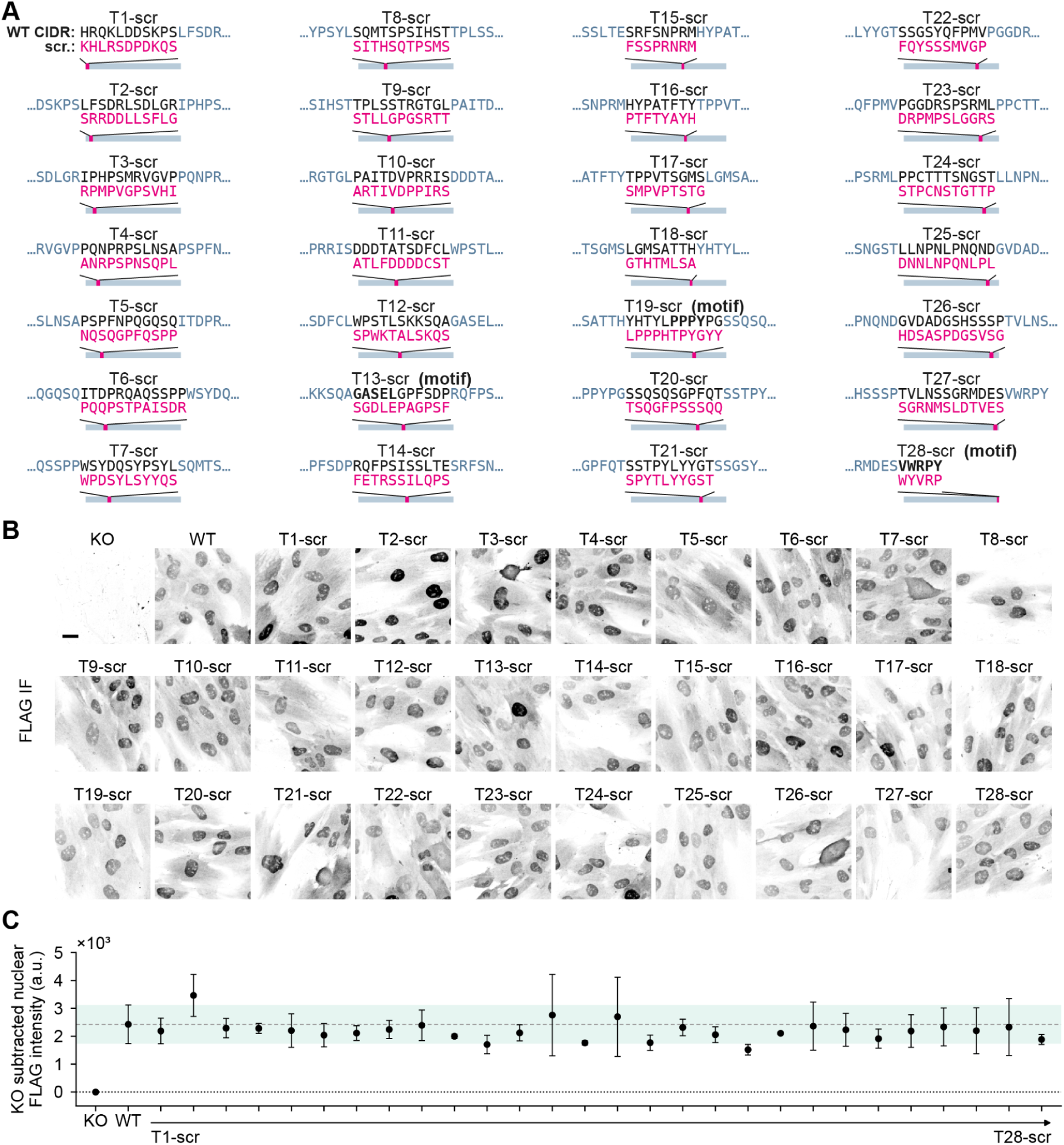
CIDR tile scramble sequences and expression. **(A)** Schematic of tile scramble constructs showing WT sequence (black) and scrambled sequence (magenta). Motif-containing tiles indicated. **(B)** Anti-FLAG immunofluorescence on day 4 of osteoblast differentiation in KO, WT and the 28 tile scramble iRUNX2 rescue lines (T1-scr–T28-scr) from Fig. 2B. Scale bar 20 µm. **(C)** KO-subtracted nuclear intensity of cells in (A), quantified as in Fig. S3B. Dots and error bars are the mean and s.d. of each clone’s mean nuclear intensity (n = 2 clones for KO). Shaded region and dashed line are WT mean and s.d.; dotted line is the KO floor. Statistics as in Fig. S3. *p < 0.05 vs. WT. No star indicates not significant vs. WT.

**Fig. S6.**
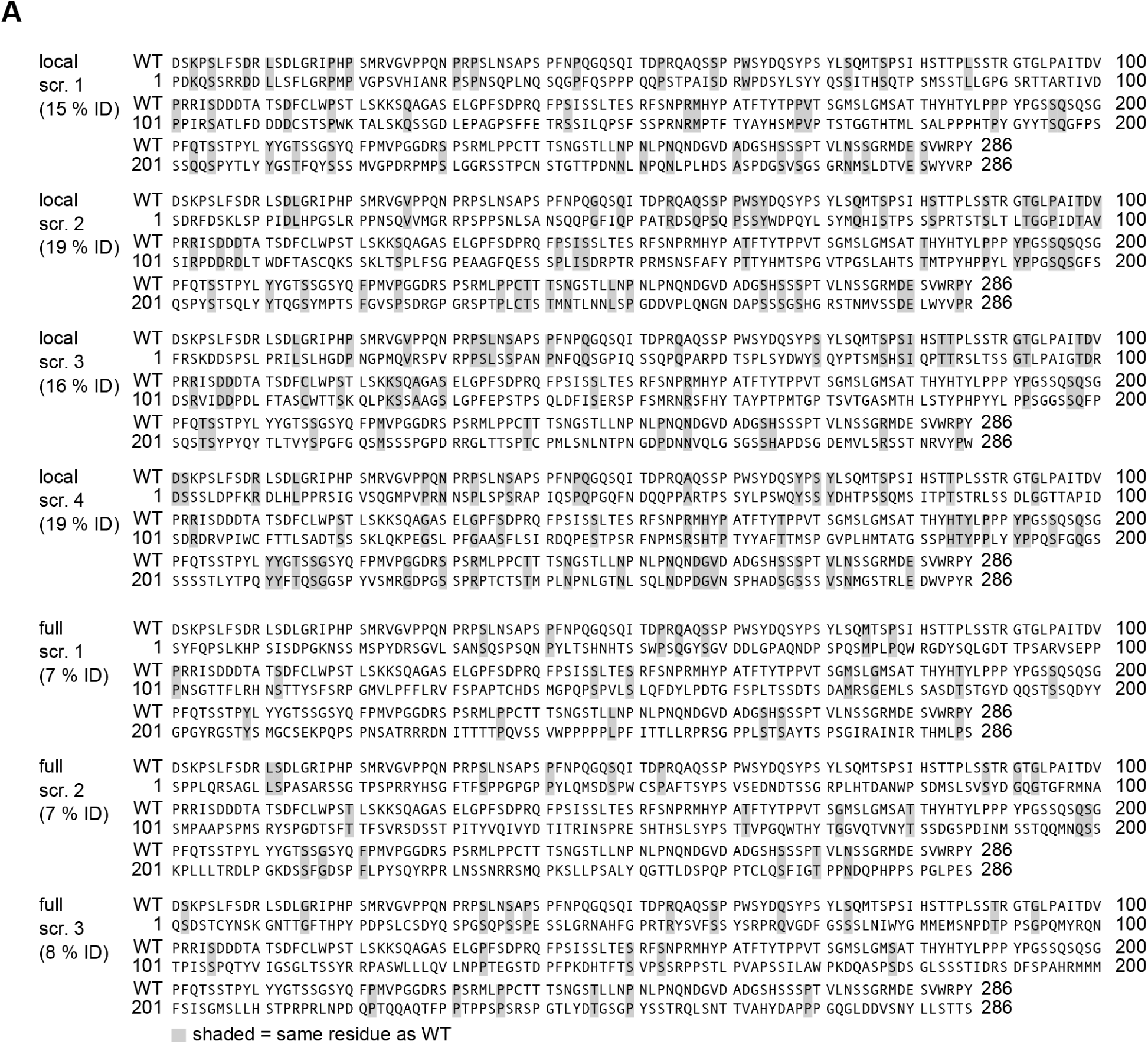
Alignment of local and full scramble variants to WT RUNX2 CIDR. **(A)** Each local scramble (LocScr1–4) and full scramble (FullScr1–3) variant was aligned to the WT RUNX2 CIDR. Grey shading marks residues identical to WT. Percent sequence identity is indicated.

**Fig. S7.**
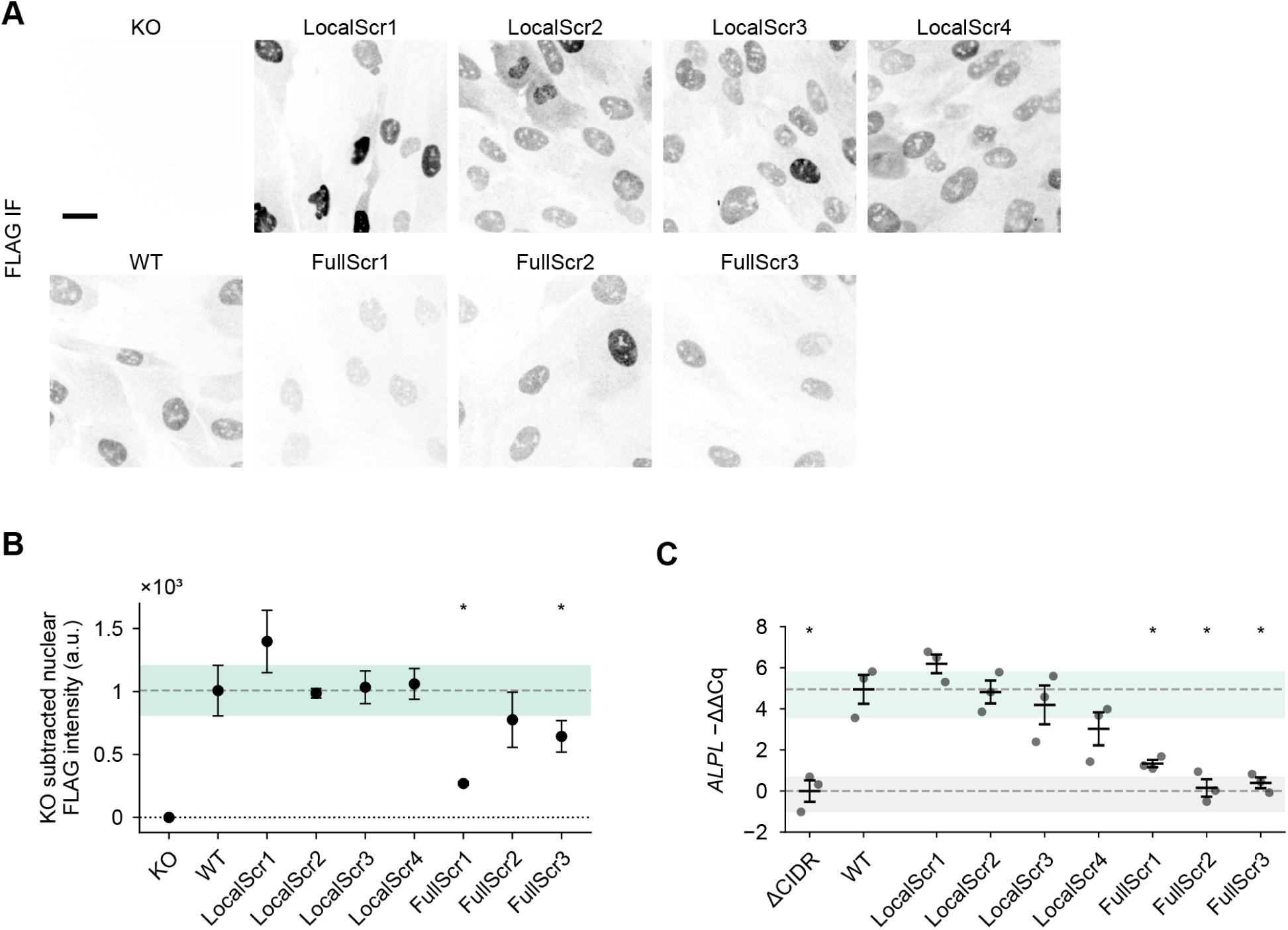
Expression and *ALPL* induction in local and full scramble CIDR variants. **(A)** Anti-FLAG immunofluorescence on day 4 of osteoblast differentiation in KO, WT, LocScr1–4 and FullScr1–3 iRUNX2 rescue lines. Scale bar 20 µm. **(B)** KO-subtracted nuclear intensity of cells in (A), quantified as in Fig. S3B. Dots and error bars are the mean and s.d. of each clone’s mean nuclear intensity (n = 6 clones for WT, pooled across two plates). Shaded region and dashed line are WT mean and s.d.; dotted line is the KO floor. **(C)** *ALPL* induction by RT-qPCR on day 4 of osteoblast differentiation, as −ΔΔCq relative to ΔCIDR measured on the same plate (ΔCq = Cq*_ALPL_* − Cq*_TBP_*; higher values indicate more *ALPL*); ΔCIDR and WT shown are from the plate carrying LocScr2–4. *TBP* was used as the reference gene. Grey dots are clones (n = 3 per variant). Error bars are mean and SEM of clones. Green and grey shaded regions are the WT and ΔCIDR clone ranges, with dashed lines at their means. Statistics as in Fig. S3. *p < 0.05 vs. WT. No star indicates not significant vs. WT.

**Fig. S8.**
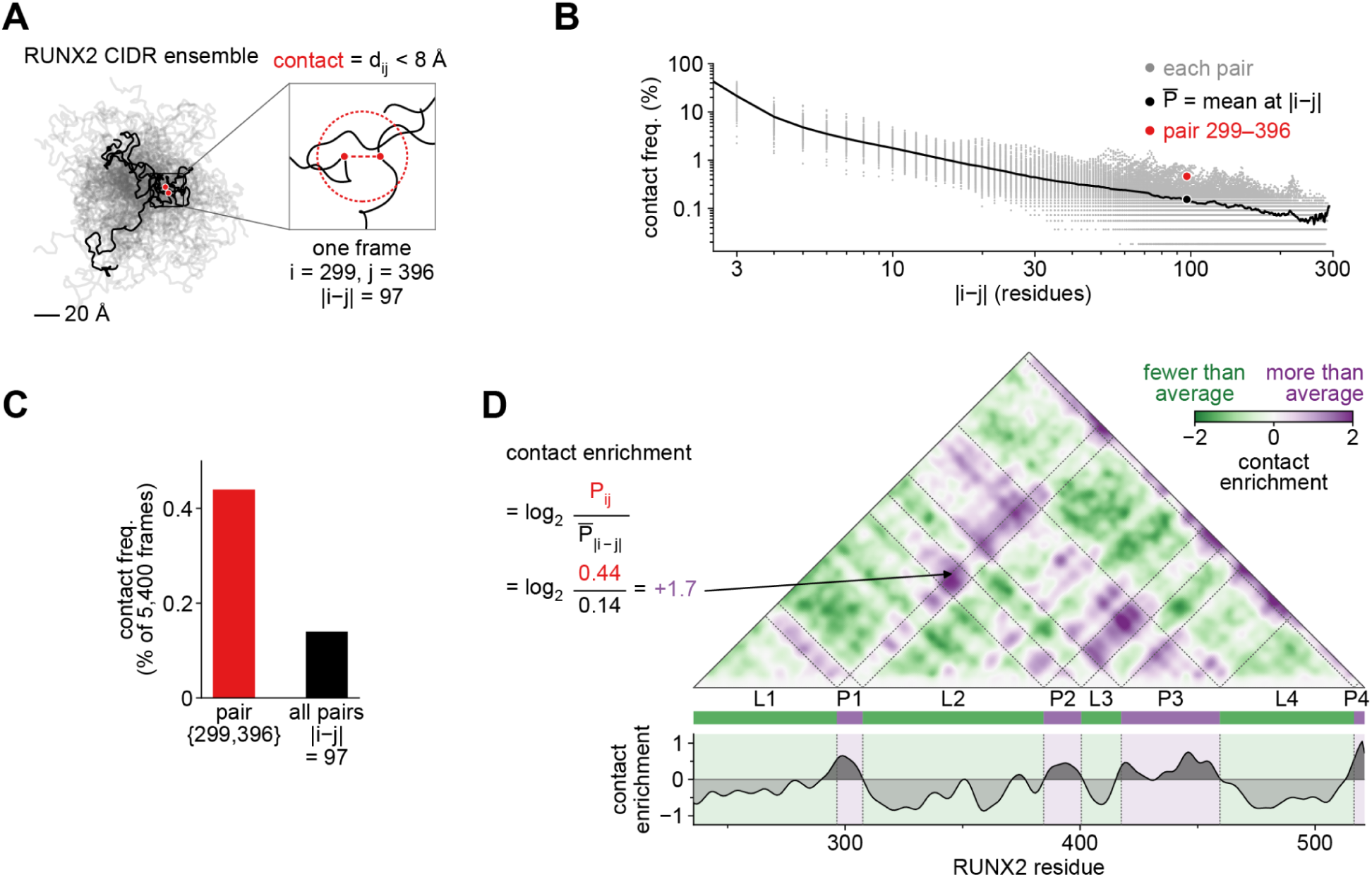
Contact frequency analysis of RUNX2 CIDR molecular dynamics simulations. (**A**) Conformers from Mpipi simulations of the WT RUNX2 CIDR (two replicates, 2,700 frames each). Scale bar, 20 Å. Inset, an example residue pair (i = 299, j = 396; |i−j| = 97) scored as in contact when beads are within 8 Å. (**B**) Contact frequency versus sequence separation for every residue pair (grey), the mean at each separation (P, black), and the example pair from A (red). (**C**) Contact frequency of the example pair (red) compared with the mean of all pairs at the same sequence separation (black). (**D**) Left, contact enrichment is log2 of a pair’s contact frequency over the mean at its sequence separation. Right, contact enrichment map for the WT CIDR with attractive patches (P1–P4) and repulsive linkers (L1–L4) annotated. Below, the per-residue enrichment track shown in Fig. 3B (three-residue moving average).

**Fig. S9.**
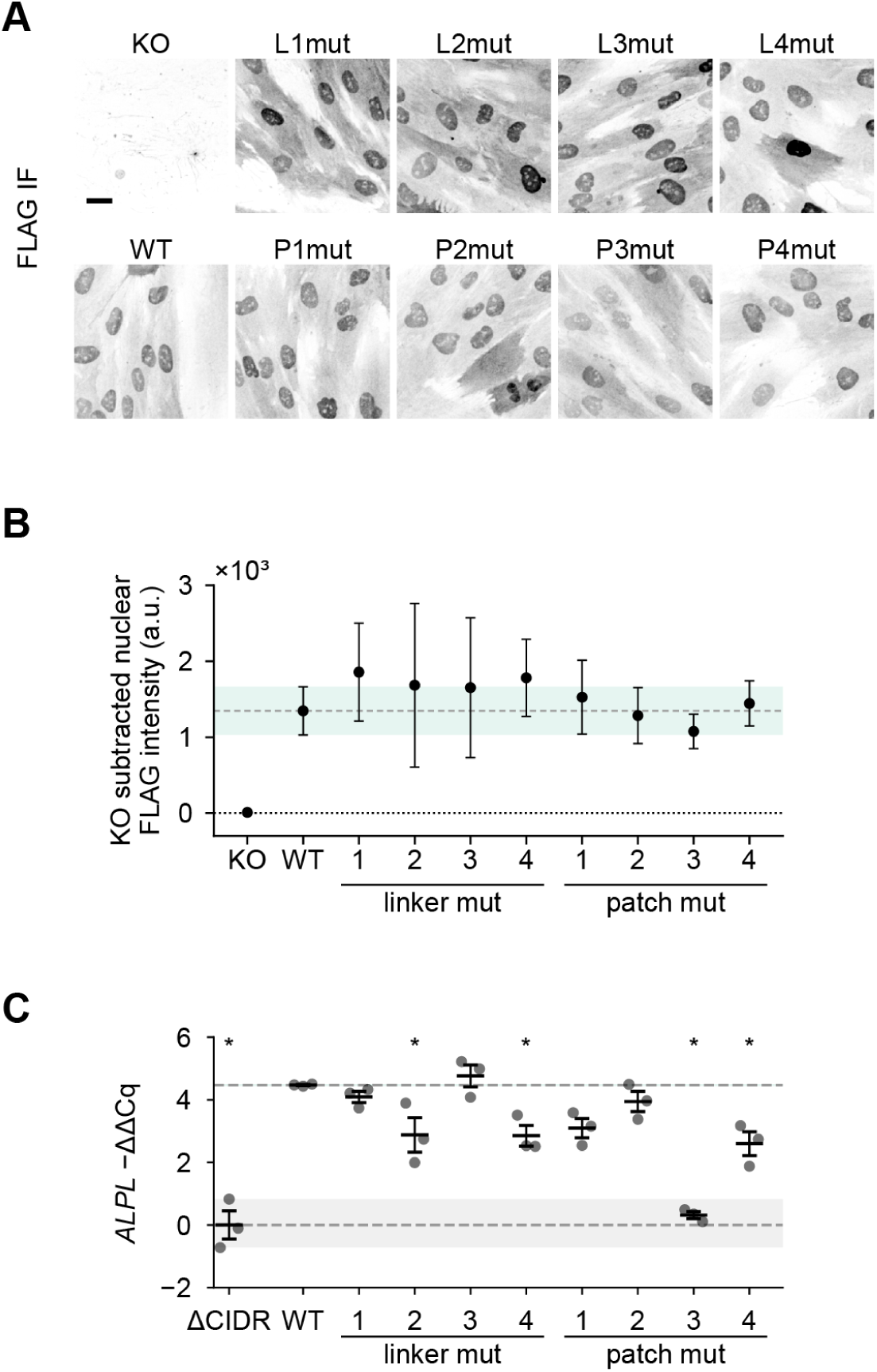
Expression and *ALPL* induction in linker and patch CIDR mutants. **(A)** Anti-FLAG immunofluorescence on day 4 of osteoblast differentiation in KO, WT, linker mutants (L1mut–L4mut) and patch mutants (P1mut–P4mut). Scale bar 20 µm. **(B)** KO-subtracted nuclear intensity of cells in (A), quantified as in Fig. S3B. Dots and error bars are the mean and s.d. of each clone’s mean nuclear intensity. Shaded region and dashed line are WT mean and s.d.; dotted line is the KO floor. **(C)** *ALPL* induction by RT-qPCR on day 4 of osteoblast differentiation, as −ΔΔCq relative to ΔCIDR (ΔCq = Cq*_ALPL_*− Cq*_TBP_*; higher values indicate more *ALPL*). *TBP* was used as the reference gene. Dots are clones (n = 3 per variant). Error bars are mean and SEM of clones. Grey shaded region is the ΔCIDR clone range; dashed line is the WT mean. Statistics as in Fig. S3. *p < 0.05 vs. WT. No star indicates not significant vs. WT.

**Fig. S10.**
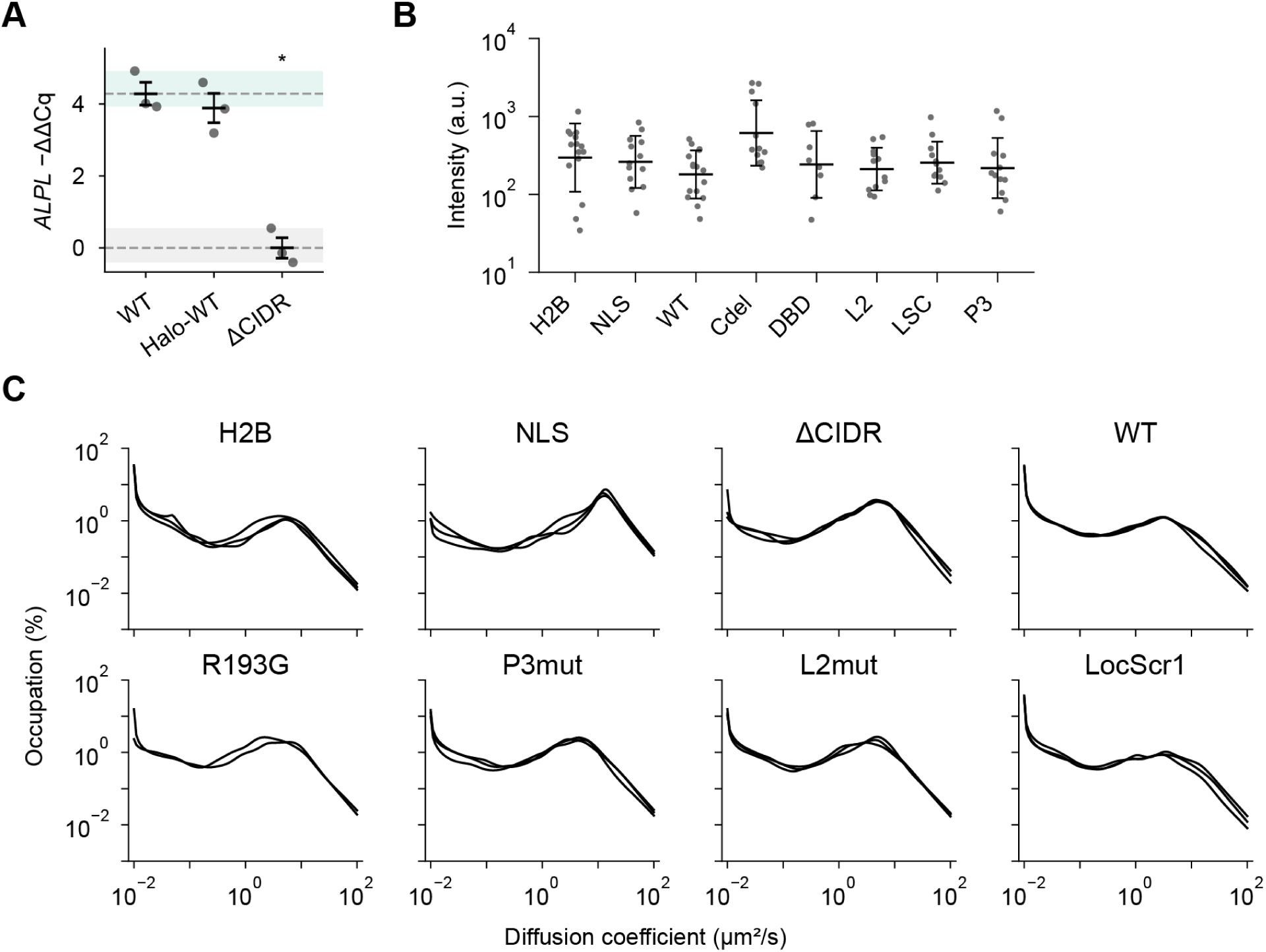
Expression and diffusion spectra from SMT experiments. **(A)** *ALPL* induction by RT-qPCR in KO cells rescued with WT, Halo-WT or ΔCIDR, shown as −ΔΔCq relative to ΔCIDR. *TBP* was used as the reference gene. Dots are clones (n = 3 per variant). Error bars are mean and SEM of clones. Green and grey shaded regions are the WT and ΔCIDR clone ranges, with dashed lines at their means. **(B)** Bulk expression from SMT experiments measured as nuclear JFX650 intensity. Dots are individual cells. Lines and error bars are mean and s.d. **(C)** saSPT occupancy spectra of diffusion coefficients for each line in (A). Each curve is the mean of per-cell posteriors from one of three independent experiments. Statistics as in Fig. S3. *p < 0.05 vs. WT. No star indicates not significant vs. WT.

**Fig. S11.**
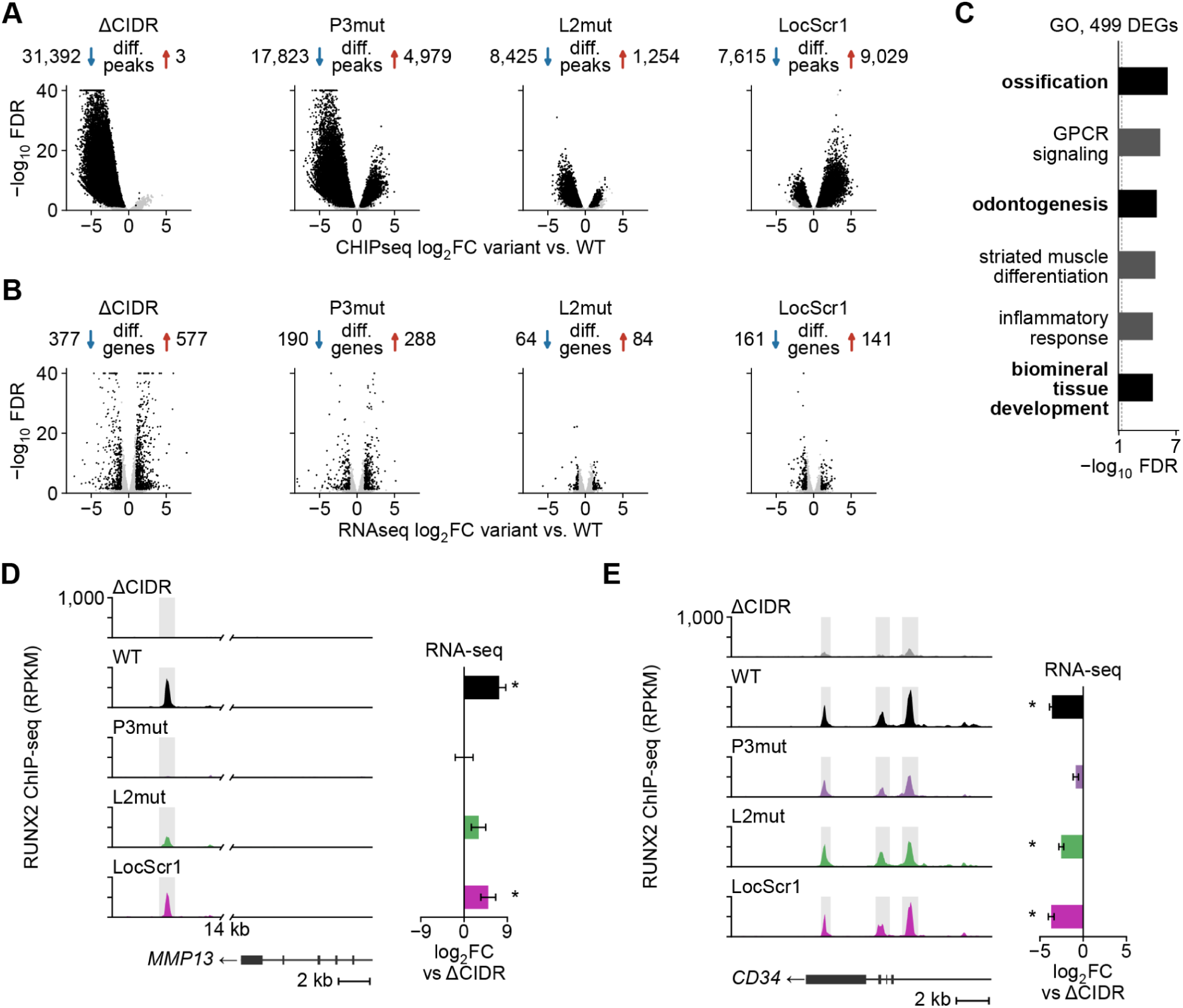
Differential binding and differential gene expression in RUNX2 CIDR variants. **(A)** RUNX2 ChIP-seq, each CIDR variant vs WT. Black, differentially bound peaks (FDR < 0.05); grey, other peaks. Peaks with less (blue) or more (red) binding in each variant shown above. **(B)** RNA-seq, each variant vs WT. Black, differentially expressed genes (FDR < 0.05, |log₂FC| ≥ 1); counts as in (A). **(C)** GO biological process terms enriched among genes differentially expressed in any variant at FDR < 0.01 and |log₂FC| ≥ 1.5 (n=499 genes). Six most significant terms; dashed line, FDR = 0.05; bold, skeletal terms. **(D, E)** RUNX2 ChIP-seq (RPKM) at peaks lost in ΔCIDR near MMP13 (C) and within CD34 (D), which are osteoblast and stem cell markers, respectively. Gene model and scale bar below. Right, RNA-seq log₂FC vs ΔCIDR ± standard error; *, differentially expressed vs ΔCIDR.

**Fig. S12.**
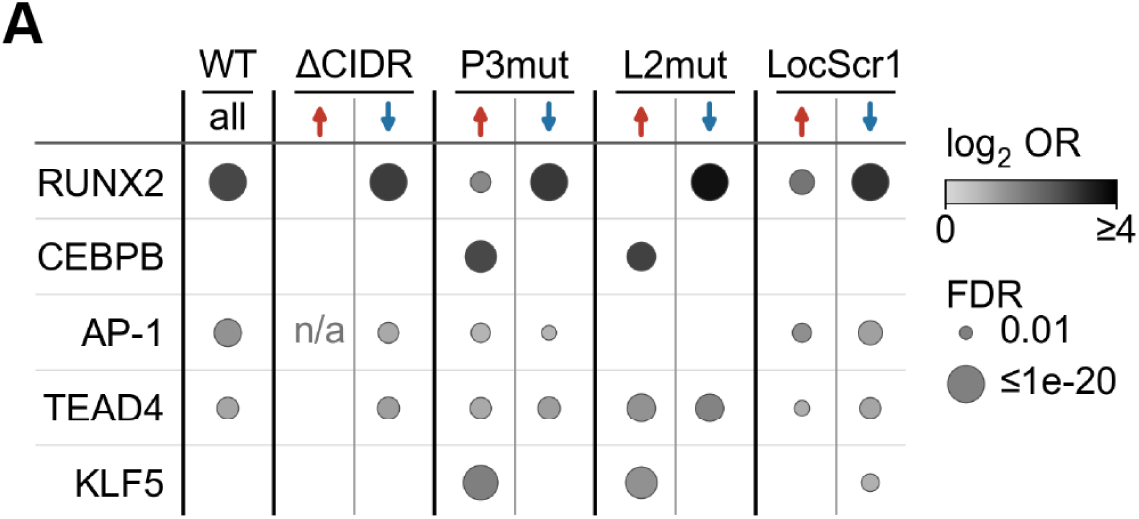
Motif enrichment in differentially bound RUNX2 CIDR CHIPseq peaks. **(A)** Enrichment of transcription factor motifs in ChIP-seq peaks that gain (red up arrow) or lose (blue down arrow) binding in each CIDR variant vs WT. The WT column shows enrichment across all reference peaks. Circle shading indicates log₂ odds ratio relative to background; circle area indicates FDR. Only motifs passing FDR < 0.01 are plotted; absent circles indicate no significant enrichment. n/a, insufficient peaks in this category. Enrichment was tested per motif by two-sided Fisher’s exact test of motif presence per set vs GC- and accessibility-matched genomic background windows; p-values were Benjamini–Hochberg corrected across all motif–set comparisons (see Methods).

**Fig. S13.**
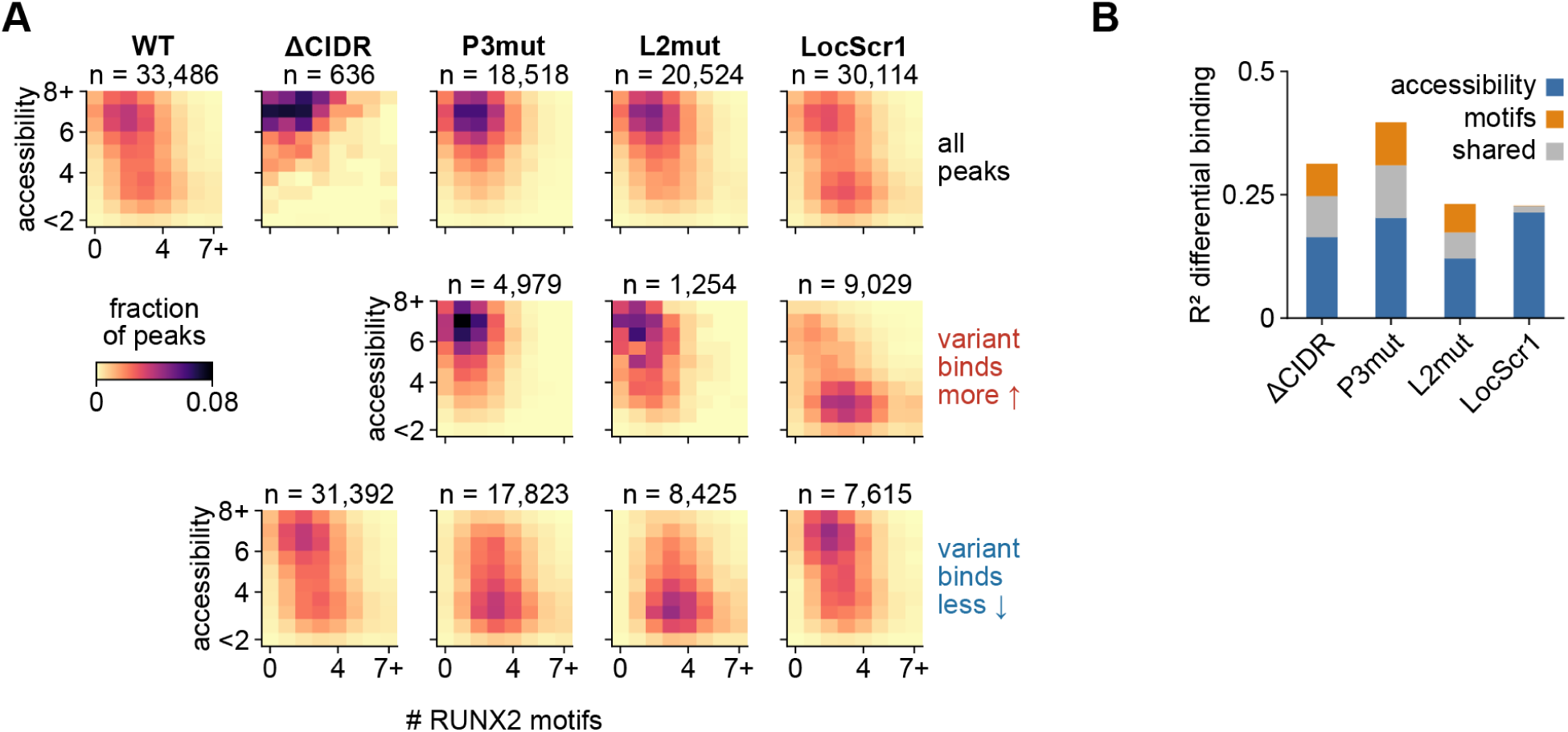
Chromatin accessibility and RUNX2 motif count at peaks differentially bound by RUNX2 CIDR variants. **(A)** Joint distribution of chromatin accessibility (DNase-seq in undifferentiated hMSCs, as in Fig. 4I) and RUNX2 motif count (FIMO) for all peaks bound by each variant (top row) and for peaks that gain (middle row, red up arrow) or lose (bottom row, blue down arrow) binding in each variant vs WT. Colour indicates the fraction of peaks in each bin; n is the number of peaks in each map. **(B)** Variance in differential binding (log₂ fold change vs WT across peaks that gain or lose binding in each variant) explained by a linear model with accessibility and RUNX2 motif count as predictors. R^2^ is partitioned into the part unique to accessibility (blue), unique to motif count (orange), and shared between them (grey).

**Fig. S14.**
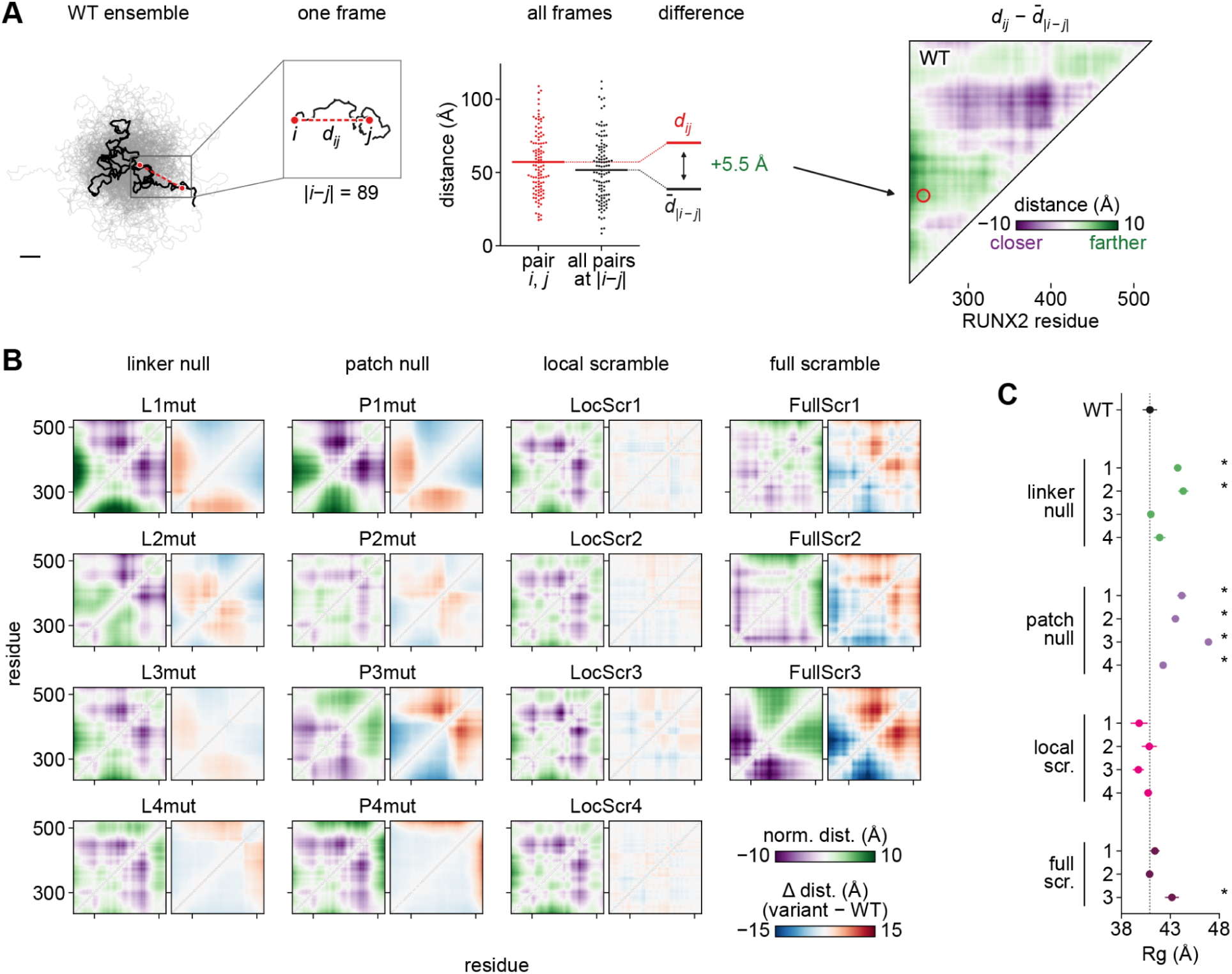
Inter-residue distance maps and chain dimensions for linker, patch, and scramble variants. **(A)** Construction of a normalized inter-residue distance map, shown for the WT CIDR. Left, 70 conformers from one Mpipi simulation (WT ensemble) with one residue pair i, j (residues 246 and 335, |i−j| = 89) marked on a single frame; scale bar, 20 Å. Middle, the distance between the pair in each frame compared with the distances of all residue pairs at the same sequence separation (110 randomly chosen frames per column; lines, means). The map value for the pair is the difference between the two means, here +5.5 Å (the pair is farther apart than expected for its separation). Right, the full WT map (upper triangle) with the worked cell ringed; purple, closer than the mean at that separation; green, farther. **(B)** Left, inter-residue distance maps from Mpipi simulations of each linker null (L1mut–L4mut), patch null (P1mut–P4mut), local scramble (LocScr1–4) and full scramble (FullScr1–3) variant. Distances are normalized to the mean inter-residue distance at matched linear sequence separation. Right, the variant − WT difference map. **(C)** Radius of gyration of each variant. Dots and error bars are mean and s.d. of three independent simulations. Dotted line is the WT mean. *p < 0.05 vs. WT, one-way ANOVA on per-simulation mean Rg with Dunnett’s test against WT; n = 3 simulations per variant.

**Fig. S15.**
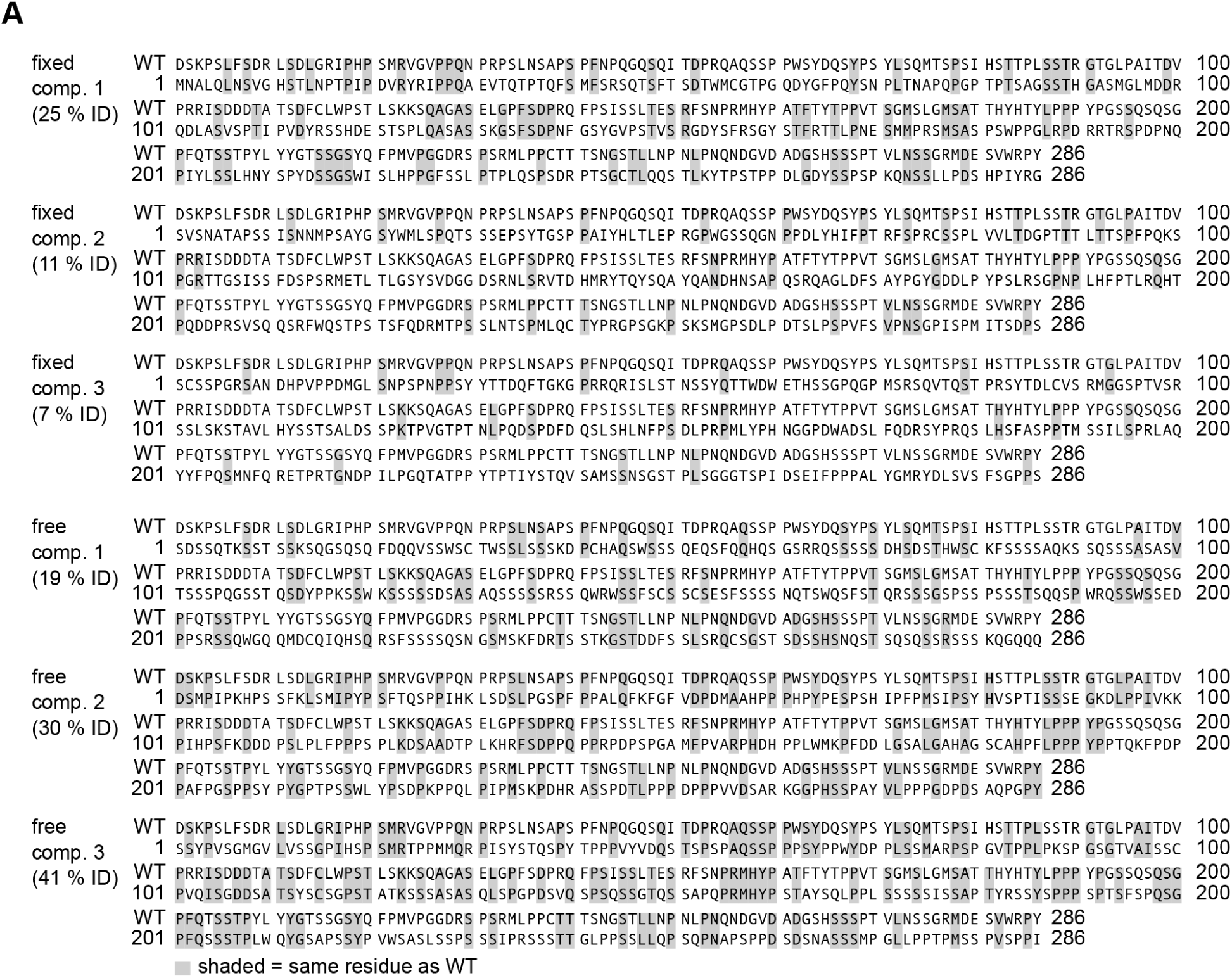
Alignment of de novo IDRs to WT CIDR. **(A)** Alignment of each fixed composition and free composition de novo designed IDR to the WT RUNX2 CIDR. Grey shading marks residues identical to WT.

**Fig. S16.**
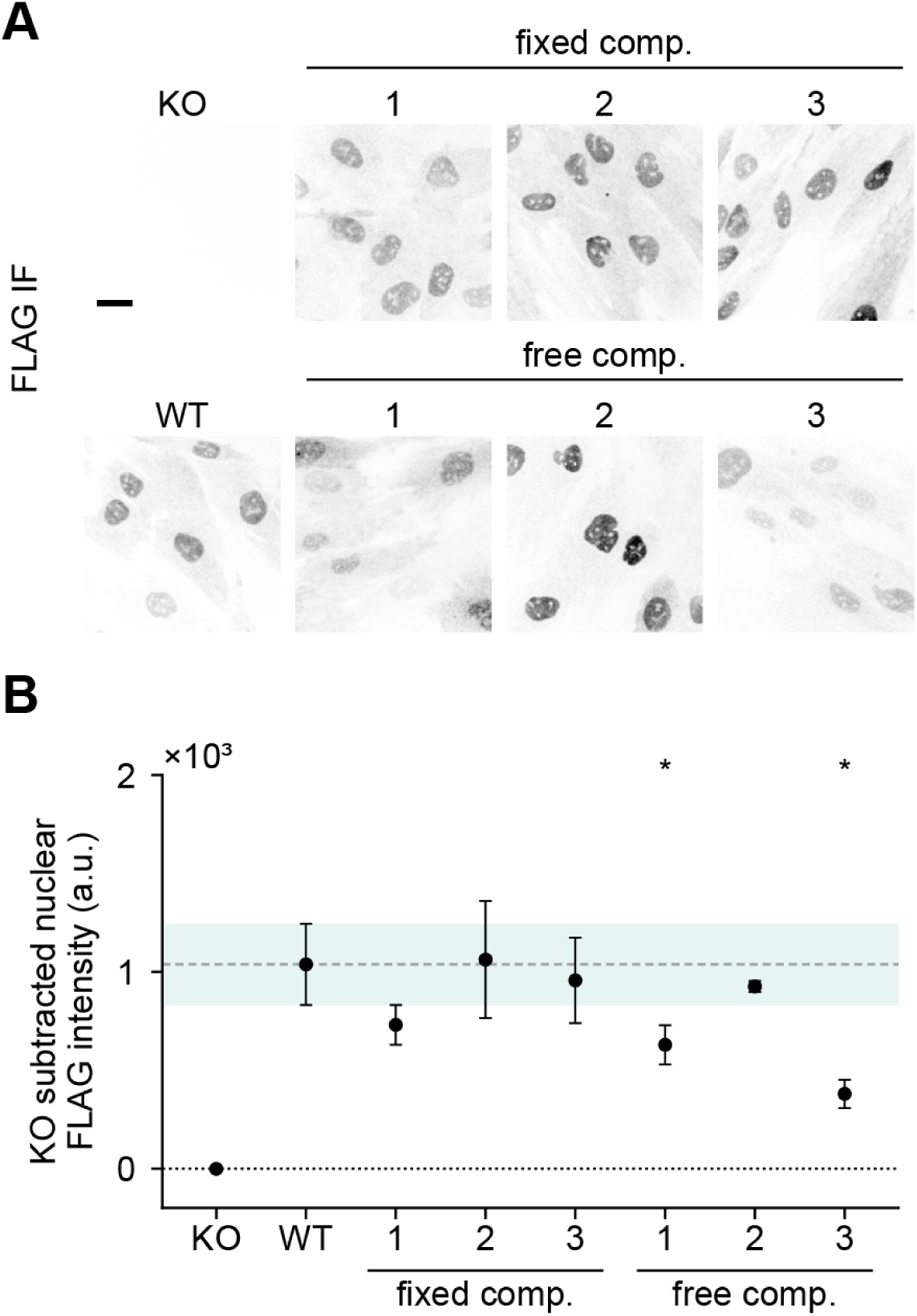
Expression of *de novo* IDR designs. **(A)** Anti-FLAG immunofluorescence on day 4 of osteoblast differentiation in KO, WT, and fixed composition (1–3) and free composition (1–3) de novo designed CIDR iRUNX2 rescue lines. Scale bar 20 µm. **(B)** KO-subtracted nuclear intensity of cells in (A), quantified as in Fig. S3B. Dots and error bars are the mean and s.d. of each clone’s mean nuclear intensity (n = 3 clones per variant). Shaded region and dashed line are WT mean and s.d.; dotted line is the KO floor. Statistics as in Fig. S3. *p < 0.05 vs. WT. No star indicates not significant vs. WT.

## Materials and methods

### Cell culture

ASC52telo (ATCC, SCRC-4000) immortalized human mesenchymal stem cells (MSCs) were maintained in DMEM (Gibco, 11-965-092) supplemented with 10% fetal bovine serum (VWR, 97068-086) and 1% penicillin–streptomycin (Gibco, 15140122) at 37 °C, 5% CO₂. Cells were passaged using TrypLE Express Enzyme (Gibco, 12563011) and were routinely tested for mycoplasma.

### Osteoblast differentiation

MSCs were seeded 2 days before inducing differentiation at 1000 cells/mL. Osteoblast differentiation media (growth media with 50 µg/mL ascorbic acid, 10 mM sodium beta-glycerophosphate, 10 nM 1α, 25-Dihydroxy-Vitamin D_3_, and 10 nM dexamethasone) was prepared fresh before the start of each experiment. Doxycycline was added to osteoblast media at 1µg/mL for the first two days to induce RUNX2 expression in all experiments involving iRUNX2 rescue cell lines. Osteoblast media was refreshed every two days for fourteen days during differentiation.

### Generation of RUNX2 knockout lines

RUNX2 knockout cells were generated using a chemically modified crRNA targeting exon 6 of RUNX2 (protospacer 5′-GTATGAGAGTAGGTGTCCCG-3′; IDT Alt-R CRISPR-Cas9 crRNA custom synthesis) using the Alt-R CRISPR-Cas9 ribonucleoprotein (RNP) system (Integrated DNA Technologies, IDT) according to the manufacturer’s instructions. RNPs were delivered to 100,000 cells using the P1 S nucleofector kit (Lonza, V4XP-1032) according to manufacturers instructions. Cells were recovered and expanded for one week and then cloned by limiting dilution in 96-well plates. Clones were verified by western blot and genomic PCR of the region surrounding the gRNA cut site (forward primer 5′-CCGAGACCAACAGAGTCAGTGAGTGCTCTCT-3′, reverse 5′-GTTTGCTCAAATATTGGGAGCTGCCGGAGTCAATAAT-3′) followed by sequencing (plasmidsaurus).

### Immunoblotting

WT and RUNX2 KO MSCs were lysed on day 4 of osteoblast differentiation in 200 μL RIPA buffer (Thermo, 89901) containing protease inhibitor tablets (Roche, 04693116001). Cells were scraped and lysate was collected and sonicated on low power at 4C for 3x pulses of 10 sec each. Lysates were then spun at 16000g for 5 min and supernatant was collected and stored at −80C. Lysates were denatured by adding NuPAGE LDS Sample Buffer (Invitrogen, NP0007) with NuPAGE Sample Reducing Agent (Invitrogen, NP0009) to 1x and incubating at 70 °C for 10 min. 20 μL of each sample was loaded onto a NuPAGE 4–12% Bis-Tris gel (Invitrogen, NP0322BOX) in NuPAGE MOPS SDS Running Buffer (Invitrogen, NP0001) and run at 200V for 30 min. Gels were transferred to nitrocellulose membranes using the TransBlot semi-dry system (BioRad, 1703940) on the mixed molecular weight setting. Membranes were blocked with 5% milk in TBST (20 mM tris-HCL, 150 mM NaC, 0.1% Tween) for 1 hr at room temp. Membranes were then stained with 1:1000 anti-RUNX2 (D1L7F, Cell Signaling, 12556T) and 1:1000 anti-TUB1A1 (Thermo, SAB4200776) in milk for 1 hr at room temp. Membranes were washed 3x in TBST and stained with 1:20,000 800CW goat anti-mouse and 1:20000 680RD goat anti-rabbit antibodies (Licor, 926-32210 and 926-68071) for 1 hr at room temp. Membranes were washed 3x in TBST and imaged on a Licor Clx imager in both 700 and 800 nm channels.

### Generation of iRUNX2 rescue lines

Plasmids containing RUNX2 variants under a doxycycline inducible protomotor and an upstream PGK promoter expression cassette containing rTTA and the puromycin resistance gene were delivered to MSCs by electroporation using a NEPA21 electrokinetic transfer system (Bulldog Bio). For all rescue experiments, RUNX2 was N-terminally FLAG tagged. For single molecule tracking experiments HaloTag was fused to the N-terminus with a glycine-serine linker. Cells were dissociated with TrypLE, washed 2x with 5 mL optimem, and 250,000 cells per sample were resuspended in 20 μL optimem. 1.5 ug of RUNX2 variant plasmid, 1 ug of piggybac transposase plasmid, and 1 μL of 10 mg/mL polyglutamic acid were added to each electroporation reaction. 20 µL of the reaction mixture was transferred to a 2mm electroporation cuvette (Bulldog bio, 12358-346) and electroporated using the settings 100V, 5 msec pulse length, 2 pulses, 10% decay rate, + polarity for the poring pulse and 20V, 50 msec pulse length, 50 msec pulse delay, 5 pulses, 40% decay rate, +/-polarity for the transfer pulse. Cells were resuspended in growth media and transferred to 6-well dishes to recover for 4 days. Puromycin was added at 0.75 µg/mL in growth media for 14 days to select for stable integrants.

### Mineralization assay

On day 14 of differentiation cells were washed 2x with PBS, and stained with 50 nM 800 CW BoneTag (Licor, 926-09375) in PBS for 10 min. Cells were washed 3x with 100 μL PBS and then imaged in 100 μL HBSS on a Licor CLx imager with focus set to 4 mm. BoneTag signal in each well was quantified using the in-cell-western western settings in the Licor software. For initial characterization of parental and KO RUNX2 clones FAM-ZOL dye was used to stain differentiated osteoblasts using the same protocol as described for BoneTag staining, and cells were imaged on a Cytation 5 multimodal plate imager using the 488 channel.

### Immunofluorescence

For all immunofluorescence experiments cells were seeded into #1.5 coverglass bottom 96-well plates (Cellvis, P961.5HN) as described for osteoblast differentiation. On day 4 of differentiation cells were fixed in 4% paraformaldehyde in PBS. Cells were permeabilized with 0.5% triton in PBS for 10 min at room temperature. Cells were then blocked in PBS containing 0.1% tween-20 and 1% bovine serum albumin (blocking solution) for 30 min at room temperature and stained with 1:1000 mouse anti-FLAG M2 (Sigma, F3165) for rescue experiments or rabbit anti-RUNX2 (D1L7F, Cell Signaling, 12556T) in blocking solution (1% BSA in PBST). Cells were washed 3x with PBST and stained with 1:5000 secondary antibody in blocking solution. Goat anti-mouse 647 (Thermo, A-21235) was used with anti-FLAG and goat anti-rabbit 647 (Thermo, A-21244) with anti-RUNX2. Cells were washed 3x with PBST and stained with 1 µg/mL Hoescht for 10 min in PBS. Cells were imaged in PBS on a Nikon Spinning disk confocal microscope using JOBS to automate multiwell plate acquisition.

### RT-qPCR

All RT-qPCR was performed using the Luna Cell Ready probe RT-qPCR kit (NEB E3031S) according to manufacturers instructions. The probes used were Hs01029144_m1 for ALPL (thermo).

### Single molecule tracking

HaloTag-RUNX2 fusion cell lines were seeded onto #1.5 coverglass bottom 96-well plates (Cellvis, P961.5HN) two days before imaging and induced with 1 ug/mL doxycycline one day before imaging. Cells were stained with 10 nM PA-JF549 dye (for single molecule tracking) and 100 nM JFX650 dye (to read out bulk RUNX2 concentration) in growth media for 20 min at 37C (*85*, *86*). Cells were washed 3x with growth media and intubated for an additional 15 min at 37C. Media was replaced with FluoroBrite (Gibco A1896701) imaging media with 10% FBS and pen/strep. SMT was performed on a custom built microscope with HILO illumination using lasers emitting at 405 nm, 561 nm, and 647 nm (*87*). Light was collected through a Nikon CFI SR HP Plan Apo 100x oil-immersion lens (NA=1.35) onto two Photometrics Prime 95B cameras. The microscope was controlled with Micro-Manager 2.0.3 (*88*). Cameras were read out in 12-bit with a 110 × 110 pixel region centred on the nucleus of each cell. Before each movie, one image of the 637 channel was taken to quantify RUNX2 expression level (20 mW laser power, 10 ms exposure). Each cell was then imaged continuously for 6,000 frames in the 561 channel (100 mW laser power, 10 ms exposure). The sample was illuminated continuously with the 405 nm laser during acquisition to photoactivate PA-JF-549. The 405 nm laser was passed through a neutral density filter and laser power was adjusted per sample to keep the number of molecules per frame low (1-5 mW). At least four cells per variant were imaged each day and three independent experiments were performed. Frames in the 561 channel were binned 2 × 2 by summing pixels, and spots were detected and linked with trackpy (*89*). Trajectories with fewer than 3 localizations were discarded. Trajectories from each cell were analysed with saspt to estimate the bound fraction and diffusion coefficient of free molecules (*90*).

### CHIPseq

#### Cell culture and differentiation

Three iRUNX2 rescue lines per FLAG-tagged variant were seeded at 1000 cells/mL into 15 cm tissue culture treated plastic dishes 2 days prior to osteoblast differentiation. Cells were induced with 1µg/mL doxycycline in osteoblast media on day 0 as described above. On day 2 cells were switched to osteoblast media without dox. On day 4 cells were harvested for CHIPseq. 5 million cells per cell line were used as input to CHIPseq.

#### Chromatin crosslinking and preparation

Chromatin immunoprecipitation and library prep was performed essentially as described (*91*). Protease inhibitors (Roche, cOmplete; 25×) were added fresh to all lysis and sonication buffers immediately before use. Cell lines were expanded in growth media in 3 15 cm dishes per line. Around 5 million cells were collected per cell line. Cells were dissociated with TrypLE, pelleted at 300 rcf for 4 min, and resuspended in 10 mL PBS. Di(N-succinimidyl) glutarate (DSG) was added to 2mM and cells were incubated for 15 min. 35 mL of PBS was added and then 5 mL of cold freshly prepared fixation solution (50 mM HEPES, 100mM NaCl, 1 mM EDTA, 0.5 mM EGTA, 11% formaldehyde) for a final concentration of 1% formaldehyde. Cells were incubated for 15 min at room temperature and the reaction was quenched with 2.5 mL of 2.5 M glycine. Cells were spun at 1100 rcf for 6 min at 4C, buffer was aspirated and the pellet was frozen at −80 for 30 min. Pellet was resuspended in lysis buffer (50 mM HEPES-KOH pH 7.5, 140 mM NaCl, 1 mM EDTA, 10% glycerol, 0.5% IGEPAL, 0.25% Triton X-100), rocked for 10 min at 4C, and centrifuged at 2300 rcf for 5 min at 4C to pellet nuclei.

#### Sonication

Supernatant was discarded and nuclei were resuspended in 2 mL sonication buffer (50 mM HEPES-KOH pH 7.5, 140 mM NaCl, 1 mM EDTA, 1 mM EGTA, 1% Triton X-100, 0.1% sodium deoxycholate, 0.1% SDS). 0.45 g sonication beads per reaction were added to sonication tubes (Diagenode, C01020031) and washed with 600 μL PBS. 1 mL of lysate was added to each sonication tube. 50 μL of lysate was kept as an input fraction. Sonication was performed in a Diagenode Bioruptor Pico sonicator using 10 cycles of 30 sec ON and 30 sec OFF. The sonicated supernatant was removed from the beads and spun at 20,000g for 10 min at 4C and the supernatant was transferred to a new tube for immunoprecipitation.

#### Immunoprecipitation

50 µL of Protein G dynabeads (Thermo, 10003D) were used per sample. Beads were washed 3x with 1 mL PBS containing 1% bovine serum albumin. Beads were resuspended in 250 µL blocking solution and 5 µg of mouse anti-FLAG M2 antibody (Sigma, F3165) were added. The antibody bead complex was rotated at 4C overnight, then washed 3x with 1 mL block solution and resuspended in 50 µL block solution. Beads were added to lysate and rotated at 4C overnight. Beads were collected on a magnet and washed once each with 1 mL Sonication Buffer, 1 mL high-salt Sonication Buffer (sonication buffer but 500 mM NaCl and 1 mM EGTA), 1 mL LiCl Buffer (20 mM Tris-HCl pH 8.0, 1 mM EDTA, 250 mM LiCl, 0.5% IGEPAL, 0.5% sodium deoxycholate) and 1 mL TE (10 mM Tris-HCl pH 8.0, 1 mM EDTA). Beads were resuspended in 210 µL Elution Buffer (50 mM Tris-HCl pH 8.0, 10 mM EDTA, 1% SDS) and incubated at 95 °C for 5 min. 200 µL of supernatant was collected and incubated overnight at 65C to reverse crosslinks. Input samples were brought to 200 μL with elution buffer and incubated in parallel.

#### DNA purification

RNase A was added to 0.2 mg/mL and samples were incubated for 2h at 37C. CaCl_2_ (10 mM final concentration) and proteinase K (0.2 mg/mL final concentration) were added and the samples were incubated at 55C for 30 min. 400 µL of phenol chloroform:isomyl alcohol were added and samples were mixed on a vortexer and spun for 5 min at 10000g at room temperature. 300 µL aqueous layer was transferred to a microfuge tube and NaCl (200 mM final concentration), 30 µg glycogen, and 880 µL of 100% ethanol were added. Samples were incubated at −80C for 1 hr, spun at 14000g for 15 min at 4C and pellets were washed with 500 μL ice cold 70% ethanol. Pellets were dried and resuspended in 70 µL water.

*Library preparation*.

NEBNext UltraExpress DNA library prep kit (NEB, E3325) was used to prep DNA according to manufacturers specifications. Libraries were run on a high sensitivity DNA tape station chip (5067–5593) for QC and to measure fragment size distribution, and concentration was quantified using the qubit high sensitivity DNA assay kit (Thermo, Q33230). Libraries were sequenced on an illumina NovaSeq using a paired-end 2 × 151 bp kit and demultiplexed yielding 41.0–50.5 M read pairs per ChIP library.

### CHIPseq data analysis

CHIPseq data was analyzed according to ENCODE guidelines (*92*). Read pairs were aligned to the UCSC hg38 assembly with BWA-MEM (*93*). Alignments were sorted and filtered for PCR/optical duplicates and MAPQ>30 using samtools (*94*). Reads overlapping the ENCODE hg38 blacklist v2 (Amemiya et al., 2019) were then removed with bedtools (*95*). Library quality was assessed with deepTools and summarized with MultiQC (*96*, *97*). The three KO ChIP BAMs were merged into a single pooled-KO control. This pooled-KO BAM was the control for all peak calling and differential binding analyses. Peaks were called for each library with MACS2 v2.2.9.1 against the pooled-KO control (*98*). Replicate reproducibility was assessed with IDR v2.0.3 for every replicate pair within a variant (18 pairs) . Input for each replicate was its top 150,000 narrowPeaks ranked by p-value. Peaks with IDR < 0.05 were kept (*99*). Differential binding was tested with Diffbind (*100*) using the DESeq2 method (*101*). A consensus peakset was built from the MACS2 narrowpeak files keeping peaks called in ≥ 2 libraries. Peaks were re-centred on summits into 401 bp windows. Each variant was compared with WT for four total contrasts.

Principal components analysis on DiffBind normalized counts of differentially bound peaks was performed using the sklearn python package (*102*). RUNX2 motif discovery was performed on 500 randomly sampled peaks from each variant using MEME. Peaks were centered on the summit +/- 15 bp. DNase-seq from human bone-marrow MSCs immortalized with hTERT (hMSC-TERT4) was obtained from GSE113253 (*49*). Reads were trimmed with Trim Galore v0.6.6 and aligned with Bowtie2 v2.5.3 to GRCh38 (*103*). Alignments were filtered to MAPQ ≥ 30 and canonical chromosomes, and duplicates were removed with Picard v2.26.10. For each RUNX2 peak, accessibility was computed as the mean DNase RPKM over the 401 bp peak window. RUNX2 motif counts per window were generated from a genome-wide FIMO scan with the WT de novo motif. Genomic background was generated from randomly sampled windows of matching GC content as the merged peakset. Motif enrichment in differentially bound peaks was tested using a FIMO scan. A random sample of peaks outside of promoters (further than 500 bp from transcription start sites) were selected and paired with random genomic windows outside all RUNX2 peaks, blacklisted regions and promoters, matched on GC content and day-0 DNase accessibility by nearest-neighbour matching. Motif families were chosen from a screen of each peak set against its matched control with SEA (MEME suite) over the full JASPAR 2024 CORE vertebrates collection, then one representative matrix per family was rescanned with FIMO. Enrichment was quantified as the log2 odds ratio of motif presence in peaks vs matched background and tested with a two-sided Fisher’s exact test. P-values were adjusted by Benjamini–Hochberg across all motif–set comparisons, and FDR < 0.01 was considered significant. For each variant, the log2 fold change of each window vs WT (from DiffBind) was regressed by ordinary least squares on RUNX motif count and DNase accessibility. Models were fit with accessibility only, motifs only, and both. Each peak was then assigned to the nearest protein-coding TSS (GENCODE v45).

### RNAseq

Cells were seeded at 1000 cells/mL in tissue culture treated 6-well plastic dishes two days before osteoblast differentiation. Osteoblast differentiation was induced as described. On day 4, cells were lysed in 200µL trizol (Thermo, 15596026) and RNA was extracted using a directzol RNA miniprep kit (Zymo, R1054) according to manufacturers instructions. RNAseq libraries were prepared using the NEBNext UltraExpress RNA library prep kit with polyA magnetic bead selection according to manufacturers instructions. Libraries were run on a high sensitivity DNA tape station chip (5067–5593) for QC and to measure fragment size distribution, and concentration was quantified using the qubit high sensitivity DNA assay kit (Thermo, Q33230) Reads were sequenced on an illumina NovaSeq 2 x 151 bp kit and demultiplexed generating 17.4–26.3 M read pairs per RNAseq library.

### RNAseq data analysis

Reads were aligned to the human genome (hg38) using STAR v2.7.11b (*104*). Gene-level read counts were obtained with featureCounts (Subread v1.6.3) on exon features grouped by gene_id, using the UCSC hg38 ensGene annotation (*105*). Differential expression was analyzed with DESeq2 v1.46.0 (R v4.5.3), with all six conditions fit jointly (18 libraries; 3 clones × 6 variants) (*101*). Genes with ≥5 counts in at least 3 samples were retained. Pairwise contrasts were extracted with the Wald test and p-values adjusted by Benjamini–Hochberg (*106*).

Differentially expressed genes were defined as adjusted p < 0.05 and |log2 fold change| ≥ 1. Principal components analysis was performed as described for CHIPseq using the differentially expressed gene set. RNA integrity of libraries was assessed from aligned reads with RSeQC read_distribution. The 5’UTR to 3’UTR tag density was used as a degradation score, which was z-scored across libraries and included as a continuous covariate in the DESeq2 model.

### Molecular dynamics simulations

#### WT RUNX2 CIDR contact map

RUNX2 CIDR simulations were run with the residue-level Mpipi force field (*61*) in LAMMPS (*107*). In this model each residue is one bead. Beads are joined by harmonic bonds (E = K(r − r₀)², K = 9.6 kcal mol⁻¹ Å⁻², r₀ = 3.81 Å). Non-bonded interactions are given by the Wang–Frenkel potential (ν = 1, μ = 2, pair-specific cutoff r_c = 3σ_ij) plus Debye–Hückel electrostatics (κ = 0.126 Å⁻¹, corresponding to 150 mM salt; relative dielectric 80; cutoff 35 Å). Each chain was placed in a cubic periodic box of 600 Å, started from a self-avoiding coil, energy-minimized and simulated at 300 K. The equations of motion were integrated with a 10 fs timestep under NVE integration with a Langevin thermostat (damping time 5 ps). Each replicate was run for 3.005 × 10⁸ steps. The first 5 × 10⁵ steps were treated as equilibration and not written out, leaving 3 µs of production per replicate. Coordinates were saved every 1 ns (3,001 frames). For every analysis, the first 10% of frames were discarded, leaving 2,701 frames per replicate. Replicates differed in their random seeds for the initial velocities and the thermostat. The wild-type CIDR (RUNX2 230–521, 292 residues) was simulated in two independent replicates. For each frame, all bead–bead distances were computed under the minimum-image convention (MDAnalysis 2.6.1) (*108*, *109*). Two residues were scored as in contact if their beads were within 8 Å. Contact probabilities P(i,j) were averaged over frames and then over replicates. To remove the generic dependence of contact probability on sequence separation, each residue’s contact enrichment was defined as c(i) = Σ_{j, |i−j| ≥ 1} [P(i,j) − B(|i−j|)], where B(s) is the mean contact probability over all residue pairs at separation s in the same ensemble. Positive values mark residues that make more contacts than average for their sequence separations, and negative values mark fewer. The track was smoothed with a centred three-residue moving average.

#### RUNX2 CIDR variant distance map correlations

Wild type and the 15 variants were simulated with Mpipi under the conditions above (300 K, 600 Å box, 150 mM, 3 µs), each in three independent replicates. For each replicate, the mean inter-bead distance ⟨d_ij⟩ was computed over the retained frames, after unwrapping the chain across periodic boundaries. The per-replicate maps were then averaged. To remove the dependence of distance on sequence separation, each map was normalized by subtracting, from every pair, the mean distance of all pairs at the same separation |i−j| in that same map. Pairs with |i−j| < 3 were excluded. After normalization, negative values (Å) mark residue pairs that are closer than typical for their separation, and positive values mark pairs that are farther apart. Each variant’s similarity to wild type was taken as the Pearson correlation coefficient between its normalized map and the wild-type map over all pairs with |i−j| ≥ 3. Difference maps (variant − wild type) were computed from the normalized maps. Map similarity was compared with log₁₀ of mineralization normalized to the wild type measured on the same plate (n = 15 variants) by Pearson correlation. Significance was assessed with a two-sided permutation test: mineralization values were reassigned to variants at random N = 100,000 times, and p = (1 + k)/(N + 1), where k is the number of permutations with |r| at least as large as the observed |r|.

### Generative IDR design

De novo CIDRs were designed using GOOSE against the FINCHES mean-field interaction model (*7*, *8*). For a candidate sequence, a self-interaction (ε) matrix was computed with the FINCHES Mpipi frontend as the intermolecular IDR matrix of the sequence against itself, treating the whole sequence as disordered, and similarity to wild type was scored as the Pearson correlation between the candidate and wild-type matrices. All designs were 286 residues, matching the wild-type CIDR (RUNX2 236–521). The two design classes were produced by different procedures. Fixed composition designs were generated with the GOOSE sequence design framework (SequenceOptimizer) run in shuffle-only mode (just_shuffle, shuffle interval 1, five shuffles per step) (*7*). This strategy preserves exact amino acid composition and only changes residue order. The run was warm-started from a randomly shuffled WT CIDR sequence, and combined three objectives: MatchSequenceDisorder against wild type (non-exact, requiring the design to be at least as disordered as wild type, weight 20); MatchingResidues with a target of less than 114 positions, i.e. 40% positional identity to wild type (weight 1); and a custom property (SelfEpsilonByMatrix) matching the candidate’s Mpipi ε matrix to that of wild type, computed as the intermolecular IDR matrix of the sequence against itself with both disorder masks disabled (target value 0, weight 1.0, model mpipi). The optimizer was run for up to 5000 iterations (tolerance 0.01). Free composition designs were generated with GOOSE using the same objectives but without shuffle-only mode, thus not preserving amino acid composition.

### Transcription factor disorder and effector-domain predictions

IDRs in the TFome and proteome were identified with metapredict (defined as contiguous disordered stretches of at least 20 residues) (*30*, *110*). Four independent models were used for activation domain prediction on TF IDRs according to each models’ published configuration.

PADDLE (*20*) was run in the published full configuration, and a TF was called if it carried at least five consecutive 53-residue tiles with Z > 4. ADpred (*33*) was called at a probability of at least 0.8 over at least five contiguous residues. TADA (*111*) was called on a run of at least two windows scoring above 0.5. Acidic exposure (*75*) was called on 39-residue tiles with a net charge between −13 and −8 and at least six tryptophan, phenylalanine or leucine residues. Gene Ontology enrichment was computed for the 885 TFs called by no predictor family, against a background of all 1,619 TFs scored by all four families. Annotations were taken from the human GOA annotation file and the go-basic ontology (*112*). Each term was tested by a two-sided Fisher’s exact test; terms annotating fewer than eight genes in the background were dropped for lack of power, and p-values were corrected across terms by the Benjamini-Hochberg procedure (*106*), with q < 0.05 taken as significant. Measured effector-domain activity was taken from four published high-throughput screens, treated as eight arms: DelRosso et al. (2023) (*35*), minCMV for activation and pEF and PGK for repression, in K562; Tycko et al. (2020) (*34*), minCMV for activation and pEF for repression; Tycko et al. (2025) (*36*), CD2 for activation and CD43 for repression, in K562; and Alerasool et al. (2022) (*113*), a GAL4-UAS assay in HEK293, activation only. The authors’ own published calls were used from their supplementary tables. For the per-TF grid, literature effector-domain annotations were taken from the compendium of Soto et al. (2022), 924 domains across 594 TFs. Examples of TFs from different functional categories are shown.

## Notes

### Summary of Updates

Text updated and a few errors corrected.

## References

1. K. I. Albanese, S. Barbe, S. Tagami, D. N. Woolfson, T. Schiex, Computational protein design. Nat. Rev. Methods Primers 5, 13 (2025).

2. W. Yang, S. Wang, G. R. Lee, J. Z. Zhang, A. Courbet, D. Juergens, X. Wang, T. Schlichthaerle, M. Abedi, R. Ragotte, L. An, I. Kalvet, S. Pellock, L. Mihaljevic, C. Glasscock, A. Pillai, A. Broerman, N. Ennist, E. Haefner, N. McNamara-Bordewick, I. Haydon, L. Stewart, G. Bhardwaj, D. Baker, The past, present and future of de novo protein design. Nature 652, 1139–1152 (2026).

3. A. S. Holehouse, B. B. Kragelund, The molecular basis for cellular function of intrinsically disordered protein regions. Nat. Rev. Mol. Cell Biol. 25, 187–211 (2024).

4. G. Tesei, F. Pesce, K. Lindorff-Larsen, Computational design of intrinsically disordered proteins. Curr. Opin. Struct. Biol. 96, 103210 (2026).

5. R. K. Krueger, M. P. Brenner, K. Shrinivas, Generalized design of sequence-ensemble-function relationships for intrinsically disordered proteins. Nat. Comput. Sci. 6, 512–523 (2026).

6. B. Novak, J. M. Lotthammer, R. J. Emenecker, A. S. Holehouse, Accurate predictions of disordered protein ensembles with STARLING. Nature, 1–11 (2026).

7. K. Hunter, T. Brandt, K. Guadalupe, K. C. Kolamunna, J. M. Lotthammer, N. M. Shamoon, J. K. Niblo, B. Nicholson, L. M. Day, A. Martinez, A. S. Holehouse, S. Sukenik, R. J. Emenecker, Rational design of disordered proteins for sequence-function investigation. Nature 658, 250–259 (2026).

8. G. M. Ginell, R. J. Emenecker, J. M. Lotthammer, A. T. Keeley, S. P. Plassmeyer, N. Razo, E. T. Usher, J. F. Pelham, A. S. Holehouse, Sequence-based prediction of intermolecular interactions driven by disordered regions. Science 388, eadq8381 (2025).

9. K. M. Ruff, M. R. King, A. W. Ying, V. Liu, A. Pant, W. E. Lieberman, M. K. Shinn, X. Su, C. Kadoch, R. V. Pappu, Molecular grammars of predicted intrinsically disordered regions that span the human proteome. Cell 189, 323–342.e17 (2026).

10. N. E. Davey, L. Simonetti, Y. Ivarsson, The next wave of interactomics: Mapping the SLiM-based interactions of the intrinsically disordered proteome. Curr. Opin. Struct. Biol. 80, 102593 (2023).

11. M. Kumar, S. Michael, J. Alvarado-Valverde, B. Mészáros, H. Sámano-Sánchez, A. Zeke, L. Dobson, T. Lazar, M. Örd, A. Nagpal, N. Farahi, M. Käser, R. Kraleti, N. E. Davey, R. Pancsa, L. B. Chemes, T. J. Gibson, The Eukaryotic Linear Motif resource: 2022 release. Nucleic Acids Res. 50, D497–D508 (2022).

12. P. Madhu, C. Benz, L. Simonetti, M. J. Winters, M. S. Subbanna, J. Kliche, L. Gomez-Lucas, H. M. Kotb, I. Krystkowiak, D. Segal, W. T. P. Darling, S. Larsen-Ledet, M. Vieler, A. Zupancic, A. Konstantinou, F. Mihalic, J. K. Varga, L. Pagano, S. Lüchow, A. Kraemer, L. Ellingboe, K. Görlitz, G. L. Righetto, C. Kossmann, R. Xiong, O. Schueler-Furman, L. Halabelian, V. Santhakumar, C. Arrowsmith, S. Knapp, M. Erdelyi, A. Stein, R. Vincentelli, P. M. Pryciak, N. E. Davey, Y. Ivarsson, An Atlas of Short linear motif-mediated human protein-protein interactions, bioRxiv (2026)p. 2026.07.03.735260.

13. P. H. Kussie, S. Gorina, V. Marechal, B. Elenbaas, J. Moreau, A. J. Levine, N. P. Pavletich, Structure of the MDM2 oncoprotein bound to the p53 tumor suppressor transactivation domain. Science 274, 948–953 (1996).

14. I. Langstein-Skora, A. Schmid, F. Huth, D. Shabani, L. Spechtenhauser, M. Likhodeeva, F. Kunert, F. J. Metzner, R. J. Emenecker, M. O. Richardson, W. Aftab, M. J. Götz, S. K. Payer, N. Pietrantoni, V. Valka, S. K. Ravichandran, T. Bartke, K.-P. Hopfner, U. Gerland, P. Korber, A. S. Holehouse, Sequence and chemical specificity define the functional landscape of intrinsically disordered regions. Nat. Cell Biol. 28, 323–337 (2026).

15. J. H. Lobel, N. T. Ingolia, Deciphering disordered regions controlling mRNA decay in high-throughput. Nature 642, 805–813 (2025).

16. M. V. Staller, A. S. Holehouse, D. Swain-Lenz, R. K. Das, R. V. Pappu, B. A. Cohen, A high-throughput mutational scan of an intrinsically disordered acidic transcriptional activation domain. Cell Syst. 6, 444–455.e6 (2018).

17. P. Tompa, M. Fuxreiter, Fuzzy complexes: polymorphism and structural disorder in protein-protein interactions. Trends Biochem. Sci. 33, 2–8 (2008).

18. R. Sharma, Z. Raduly, M. Miskei, M. Fuxreiter, Fuzzy complexes: Specific binding without complete folding. FEBS Lett. 589, 2533–2542 (2015).

19. L. M. Tuttle, D. Pacheco, L. Warfield, J. Luo, J. Ranish, S. Hahn, R. E. Klevit, Gcn4-Mediator specificity is mediated by a large and dynamic fuzzy protein-protein complex. Cell Rep. 22, 3251–3264 (2018).

20. A. L. Sanborn, B. T. Yeh, J. T. Feigerle, C. V. Hao, R. J. Townshend, E. Lieberman Aiden, R. O. Dror, R. D. Kornberg, Simple biochemical features underlie transcriptional activation domain diversity and dynamic, fuzzy binding to Mediator. Elife 10, e68068 (2021).

21. H. Lyons, R. T. Veettil, P. Pradhan, C. Fornero, N. De La Cruz, K. Ito, M. Eppert, R. G. Roeder, B. R. Sabari, Functional partitioning of transcriptional regulators by patterned charge blocks. Cell 186, 327–345.e28 (2023).

22. V. Mindel, S. Brodsky, A. Cohen, W. Manadre, F. Jonas, M. Carmi, N. Barkai, Intrinsically disordered regions of the Msn2 transcription factor encode multiple functions using interwoven sequence grammars. Nucleic Acids Res. 52, 2260–2272 (2024).

23. E. W. Martin, A. S. Holehouse, I. Peran, M. Farag, J. J. Incicco, A. Bremer, C. R. Grace, A. Soranno, R. V. Pappu, T. Mittag, Valence and patterning of aromatic residues determine the phase behavior of prion-like domains. Science 367, 694–699 (2020).

24. Z. Du, H. Wang, S. Luo, Z. Yun, C. Wu, W. Yang, M. Buck, W. Zheng, A. L. Hansen, H.-Y. Kao, S. Yang, The sequence-structure-function relationship of intrinsic ERα disorder. Nature 638, 1130–1138 (2025).

25. G. Koren, S. Meir, L. Holschuh, H. D. T. Mertens, T. Ehm, N. Yahalom, A. Golombek, T. Schwartz, D. I. Svergun, O. A. Saleh, J. Dzubiella, R. Beck, Intramolecular structural heterogeneity altered by long-range contacts in an intrinsically disordered protein. Proc. Natl. Acad. Sci. U. S. A. 120, e2220180120 (2023).

26. W. Borcherds, F.-X. Theillet, A. Katzer, A. Finzel, K. M. Mishall, A. T. Powell, H. Wu, W. Manieri, C. Dieterich, P. Selenko, A. Loewer, G. W. Daughdrill, Disorder and residual helicity alter p53-Mdm2 binding affinity and signaling in cells. Nat. Chem. Biol. 10, 1000–1002 (2014).

27. N. S. González-Foutel, J. Glavina, W. M. Borcherds, M. Safranchik, S. Barrera-Vilarmau, A. Sagar, A. Estaña, A. Barozet, N. A. Garrone, G. Fernandez-Ballester, C. Blanes-Mira, I. E. Sánchez, G. de Prat-Gay, J. Cortés, P. Bernadó, R. V. Pappu, A. S. Holehouse, G. W. Daughdrill, L. B. Chemes, Conformational buffering underlies functional selection in intrinsically disordered protein regions. Nat. Struct. Mol. Biol. 29, 781–790 (2022).

28. H. J. Dyson, P. E. Wright, Equilibrium NMR studies of unfolded and partially folded proteins. Nat. Struct. Biol. 5 **Suppl**, 499–503 (1998).

29. J. Liu, N. B. Perumal, C. J. Oldfield, E. W. Su, V. N. Uversky, A. K. Dunker, Intrinsic disorder in transcription factors. Biochemistry 45, 6873–6888 (2006).

30. S. A. Lambert, A. Jolma, L. F. Campitelli, P. K. Das, Y. Yin, M. Albu, X. Chen, J. Taipale, T. R. Hughes, M. T. Weirauch, The human transcription factors. Cell 172, 650–665 (2018).

31. L. F. Soto, Z. Li, C. S. Santoso, A. Berenson, I. Ho, V. X. Shen, S. Yuan, J. I. Fuxman Bass, Compendium of human transcription factor effector domains. Mol. Cell 82, 514–526 (2022).

32. A. Jolma, K. U. Laverty, A. Fathi, A. W. H. Yang, I. Yellan, I. E. Vorontsov, A. J. Gralak, J. F. Kribelbauer-Swietek, S. Inukai, R. Razavi, M. Albu, A. Brechalov, Z. M. Patel, V. Nozdrin, G. Meshcheryakov, A. Buyan, I. Kozin, S. Abramov, A. Boytsov, Codebook Consortium, Q. Morris, M. T. Weirauch, O. Fornes, V. J. Makeev, J. Grau, I. Grosse, P. Bucher, B. Deplancke, I. V. Kulakovskiy, T. R. Hughes, An expanded codebook of human transcription factor DNA-binding specificity. Nature 657, 275–283 (2026).

33. A. Erijman, L. Kozlowski, S. Sohrabi-Jahromi, J. Fishburn, L. Warfield, J. Schreiber, W. S. Noble, J. Söding, S. Hahn, A high-throughput screen for transcription activation domains reveals their sequence features and permits prediction by deep learning. Mol. Cell 78, 890–902.e6 (2020).

34. J. Tycko, N. DelRosso, G. T. Hess, Aradhana, A. Banerjee, A. Mukund, M. V. Van, B. K. Ego, D. Yao, K. Spees, P. Suzuki, G. K. Marinov, A. Kundaje, M. C. Bassik, L. Bintu, High-throughput discovery and characterization of human transcriptional effectors. Cell 183, 2020–2035.e16 (2020).

35. N. DelRosso, J. Tycko, P. Suzuki, C. Andrews, Aradhana, A. Mukund, I. Liongson, C. Ludwig, K. Spees, P. Fordyce, M. C. Bassik, L. Bintu, Large-scale mapping and mutagenesis of human transcriptional effector domains. Nature 616, 365–372 (2023).

36. J. Tycko, M. V. Van, Aradhana, N. DelRosso, H. Ye, D. Yao, R. Valbuena, A. Vaughan-Jackson, X. Xu, C. Ludwig, K. Spees, K. Liu, M. Gu, V. Khare, A. X. Mukund, P. H. Suzuki, S. Arana, C. Zhang, P. P. Du, T. S. Ornstein, G. T. Hess, R. A. Kamber, L. S. Qi, A. S. Khalil, L. Bintu, M. C. Bassik, Development of compact transcriptional effectors using high-throughput measurements in diverse contexts. Nat. Biotechnol. 43, 1525–1538 (2025).

37. A. Jolma, J. Yan, T. Whitington, J. Toivonen, K. R. Nitta, P. Rastas, E. Morgunova, M. Enge, M. Taipale, G. Wei, K. Palin, J. M. Vaquerizas, R. Vincentelli, N. M. Luscombe, T. R. Hughes, P. Lemaire, E. Ukkonen, T. Kivioja, J. Taipale, DNA-binding specificities of human transcription factors. Cell 152, 327–339 (2013).

38. N. Alerasool, H. Leng, Z.-Y. Lin, A.-C. Gingras, M. Taipale, Identification and functional characterization of transcriptional activators in human cells. Mol. Cell 82, 677–695.e7 (2022).

39. M. M. Freund, F. J. deHaro-Arbona, S. Baloul, A. Torhorst, C. Roussos, A. J. Ruffridge, A. Q. Rashoff, R. Hazzard, P. W. Lewis, S. J. Bray, M. M. Harrison, Pioneer-factor activity requires stable chromatin occupancy mediated by both sequence-specific binding and disordered protein domains. Sci. Adv. 12 (2026).

40. S. Brodsky, T. Jana, K. Mittelman, M. Chapal, D. K. Kumar, M. Carmi, N. Barkai, Intrinsically disordered regions direct transcription factor in vivo binding specificity. Mol. Cell 79, 459–471.e4 (2020).

41. F. Jonas, Y. Navon, N. Barkai, Intrinsically disordered regions as facilitators of the transcription factor target search. Nat. Rev. Genet. 26, 424–435 (2025).

42. A. A. Abidi, C. Cattoglio, N. N. Tang, V. B. Fan, G. M. Dailey, A. D. Hay, P. Kunamaneni, D. E. Milkie, X. Darzacq, E. Betzig, R. Tjian, T. G. W. Graham, Unstructured transcription factor interactions enable emergent specificity. Science 392, eaeb6487 (2026).

43. C. J. Andrews, E. J. Costa, G. L. Janer Carattini, N. V. DelRosso, T. Fujimori, M. Shimasawa, L. Bintu, Bifunctional transcriptional effector domains control gene expression pulses in an occupancy-dependent manner, bioRxivorg (2025)p. 2025.12.10.693551.

44. C. Nüsslein-Volhard, E. Wieschaus, Mutations affecting segment number and polarity in Drosophila. Nature 287, 795–801 (1980).

45. D. M. Wellik, M. R. Capecchi, Hox10 and Hox11 genes are required to globally pattern the mammalian skeleton. Science 301, 363–367 (2003).

46. T. Komori, H. Yagi, S. Nomura, A. Yamaguchi, K. Sasaki, K. Deguchi, Y. Shimizu, R. T. Bronson, Y. H. Gao, M. Inada, M. Sato, R. Okamoto, Y. Kitamura, S. Yoshiki, T. Kishimoto, Targeted disruption of Cbfa1 results in a complete lack of bone formation owing to maturational arrest of osteoblasts. Cell 89, 755–764 (1997).

47. F. Otto, A. P. Thornell, T. Crompton, A. Denzel, K. C. Gilmour, I. R. Rosewell, G. W. Stamp, R. S. Beddington, S. Mundlos, B. R. Olsen, P. B. Selby, M. J. Owen, Cbfa1, a candidate gene for cleidocranial dysplasia syndrome, is essential for osteoblast differentiation and bone development. Cell 89, 765–771 (1997).

48. S. Mundlos, F. Otto, C. Mundlos, J. B. Mulliken, A. S. Aylsworth, S. Albright, D. Lindhout, W. G. Cole, W. Henn, J. H. Knoll, M. J. Owen, R. Mertelsmann, B. U. Zabel, B. R. Olsen, Mutations involving the transcription factor CBFA1 cause cleidocranial dysplasia. Cell 89, 773–779 (1997).

49. A. Rauch, A. K. Haakonsson, J. G. S. Madsen, M. Larsen, I. Forss, M. R. Madsen, E. L. Van Hauwaert, C. Wiwie, N. Z. Jespersen, M. Tencerova, R. Nielsen, B. D. Larsen, R. Röttger, J. Baumbach, C. Scheele, M. Kassem, S. Mandrup, Osteogenesis depends on commissioning of a network of stem cell transcription factors that act as repressors of adipogenesis. Nat. Genet. 51, 716–727 (2019).

50. M. F. Pittenger, A. M. Mackay, S. C. Beck, R. K. Jaiswal, R. Douglas, J. D. Mosca, M. A. Moorman, D. W. Simonetti, S. Craig, D. R. Marshak, Multilineage potential of adult human mesenchymal stem cells. Science 284, 143–147 (1999).

51. S. Thaweesapphithak, T. Theerapanon, K. Rattanapornsompong, N. Intarak, P. Kanpittaya, V. Trachoo, T. Porntaveetus, V. Shotelersuk, Functional consequences of C-terminal mutations in RUNX2. Sci. Rep. 13, 12202 (2023).

52. L. Zeng, J. Wei, D. Han, H. Liu, Y. Liu, N. Zhao, S. Sun, Y. Wang, H. Feng, Functional analysis of novel RUNX2 mutations in cleidocranial dysplasia. Mutagenesis 32, 437–443 (2017).

53. S. J. Triezenberg, R. C. Kingsbury, S. L. McKnight, Functional dissection of VP16, the trans-activator of herpes simplex virus immediate early gene expression. Genes Dev. 2, 718–729 (1988).

54. W. C. Yeh, Z. Cao, M. Classon, S. L. McKnight, Cascade regulation of terminal adipocyte differentiation by three members of the C/EBP family of leucine zipper proteins. Genes Dev. 9, 168–181 (1995).

55. H. Akiyama, M.-C. Chaboissier, J. F. Martin, A. Schedl, B. de Crombrugghe, The transcription factor Sox9 has essential roles in successive steps of the chondrocyte differentiation pathway and is required for expression of Sox5 and Sox6. Genes Dev. 16, 2813–2828 (2002).

56. F. Jonas, M. Carmi, B. Krupkin, J. Steinberger, S. Brodsky, T. Jana, N. Barkai, The molecular grammar of protein disorder guiding genome-binding locations. Nucleic Acids Res. 51, 4831–4844 (2023).

57. S. K. Zaidi, A. J. Sullivan, R. Medina, Y. Ito, A. J. van Wijnen, J. L. Stein, J. B. Lian, G. S. Stein, Tyrosine phosphorylation controls Runx2-mediated subnuclear targeting of YAP to repress transcription. EMBO J. 23, 790–799 (2004).

58. K. Thirunavukkarasu, M. Mahajan, K. W. McLarren, S. Stifani, G. Karsenty, Two domains unique to osteoblast-specific transcription factor Osf2/Cbfa1 contribute to its transactivation function and its inability to heterodimerize with Cbfbeta. Mol. Cell. Biol. 18, 4197–4208 (1998).

59. G. L. Dignon, W. Zheng, Y. C. Kim, R. B. Best, J. Mittal, Sequence determinants of protein phase behavior from a coarse-grained model. PLoS Comput. Biol. 14, e1005941 (2018).

60. G. Tesei, T. K. Schulze, R. Crehuet, K. Lindorff-Larsen, Accurate model of liquid-liquid phase behavior of intrinsically disordered proteins from optimization of single-chain properties. Proc. Natl. Acad. Sci. U. S. A. 118, e2111696118 (2021).

61. J. A. Joseph, A. Reinhardt, A. Aguirre, P. Y. Chew, K. O. Russell, J. R. Espinosa, A. Garaizar, R. Collepardo-Guevara, Physics-driven coarse-grained model for biomolecular phase separation with near-quantitative accuracy. Nat. Comput. Sci. 1, 732–743 (2021).

62. S. Li, S. Barethiya, J. Chen, HyRes: Accurate physics-based simulation of dynamic protein structures and interactions in complex environments at scale, bioRxivorg (2026)p. 2026.06.23.734133.

63. G. L. Dignon, W. Zheng, R. B. Best, Y. C. Kim, J. Mittal, Relation between single-molecule properties and phase behavior of intrinsically disordered proteins. Proc. Natl. Acad. Sci. U. S. A. 115, 9929–9934 (2018).

64. G. Tesei, A. I. Trolle, N. Jonsson, J. Betz, F. E. Knudsen, F. Pesce, K. E. Johansson, K. Lindorff-Larsen, Conformational ensembles of the human intrinsically disordered proteome. Nature 626, 897–904 (2024).

65. N. Hess, J. A. Joseph, Biomolecular condensates dictate the folding landscape of protein alpha-helices. J. Am. Chem. Soc. 148, 36392–36413 (2026).

66. T. Lionnet, C. Wu, Single-molecule tracking of transcription protein dynamics in living cells: seeing is believing, but what are we seeing? Curr. Opin. Genet. Dev. 67, 94–102 (2021).

67. D. Ovek Baydar, I. Rauluseviciute, D. R. Aronsen, R. Blanc-Mathieu, I. Bonthuis, H. de Beukelaer, K. Ferenc, A. Jegou, V. Kumar, R. B. Lemma, J. Lucas, M. Pochon, C. M. Yun, V. Ramalingam, S. S. Deshpande, A. Patel, G. K. Marinov, A. T. Wang, A. Aguirre, J. A. Castro-Mondragon, D. Baranasic, J. Chèneby, S. Gundersen, M. Johansen, A. Khan, M. L. Kuijjer, E. Hovig, B. Lenhard, A. Sandelin, K. Vandepoele, W. W. Wasserman, F. Parcy, A. Kundaje, A. Mathelier, JASPAR 2026: expansion of transcription factor binding profiles and integration of deep learning models. Nucleic Acids Res. 54, D184–D193 (2026).

68. M. van der Deen, J. Akech, D. Lapointe, S. Gupta, D. W. Young, M. A. Montecino, M. Galindo, J. B. Lian, J. L. Stein, G. S. Stein, A. J. van Wijnen, Genomic promoter occupancy of runt-related transcription factor RUNX2 in Osteosarcoma cells identifies genes involved in cell adhesion and motility. J. Biol. Chem. 287, 4503–4517 (2012).

69. J. M. Lotthammer, G. M. Ginell, D. Griffith, R. J. Emenecker, A. S. Holehouse, Direct prediction of intrinsically disordered protein conformational properties from sequence. Nat. Methods 21, 465–476 (2024).

70. S. Müller-Späth, A. Soranno, V. Hirschfeld, H. Hofmann, S. Rüegger, L. Reymond, D. Nettels, B. Schuler, From the Cover: Charge interactions can dominate the dimensions of intrinsically disordered proteins. Proc. Natl. Acad. Sci. U. S. A. 107, 14609–14614 (2010).

71. A. H. Mao, S. L. Crick, A. Vitalis, C. L. Chicoine, R. V. Pappu, Net charge per residue modulates conformational ensembles of intrinsically disordered proteins. Proc. Natl. Acad. Sci. U. S. A. 107, 8183–8188 (2010).

72. J. A. Marsh, J. D. Forman-Kay, Sequence determinants of compaction in intrinsically disordered proteins. Biophys. J. 98, 2383–2390 (2010).

73. R. K. Das, R. V. Pappu, Conformations of intrinsically disordered proteins are influenced by linear sequence distributions of oppositely charged residues. Proc. Natl. Acad. Sci. U. S. A. 110, 13392–13397 (2013).

74. J. L. Watson, D. Juergens, N. R. Bennett, B. L. Trippe, J. Yim, H. E. Eisenach, W. Ahern, A. J. Borst, R. J. Ragotte, L. F. Milles, B. I. M. Wicky, N. Hanikel, S. J. Pellock, A. Courbet, W. Sheffler, J. Wang, P. Venkatesh, I. Sappington, S. V. Torres, A. Lauko, V. De Bortoli, E. Mathieu, S. Ovchinnikov, R. Barzilay, T. S. Jaakkola, F. DiMaio, M. Baek, D. Baker, De novo design of protein structure and function with RFdiffusion. Nature 620, 1089–1100 (2023).

75. S. R. Kotha, M. V. Staller, Clusters of acidic and hydrophobic residues can predict acidic transcriptional activation domains from protein sequence. Genetics 225, iyad131 (2023).

76. C. N. Ravarani, T. Y. Erkina, G. De Baets, D. C. Dudman, A. M. Erkine, M. M. Babu, High-throughput discovery of functional disordered regions: investigation of transactivation domains. Mol. Syst. Biol. 14, e8190 (2018).

77. P. B. Sigler, Transcriptional activation. Acid blobs and negative noodles. Nature 333, 210–212 (1988).

78. N. Ferrari, A. I. Riggio, S. Mason, L. McDonald, A. King, T. Higgins, I. Rosewell, J. C. Neil, M. J. Smalley, O. J. Sansom, J. Morris, E. R. Cameron, K. Blyth, Runx2 contributes to the regenerative potential of the mammary epithelium. Sci. Rep. 5, 15658 (2015).

79. K. Nagata, H. Hojo, S. H. Chang, H. Okada, F. Yano, R. Chijimatsu, Y. Omata, D. Mori, Y. Makii, M. Kawata, T. Kaneko, Y. Iwanaga, H. Nakamoto, Y. Maenohara, N. Tachibana, H. Ishikura, J. Higuchi, Y. Taniguchi, S. Ohba, U.-I. Chung, S. Tanaka, T. Saito, Runx2 and Runx3 differentially regulate articular chondrocytes during surgically induced osteoarthritis development. Nat. Commun. 13, 6187 (2022).

80. G. A. Meaker, M. Nicholls, C. Chahrour, I. Hsu, A. Smith, Y. Bozhilov, M. Leung, H. Vassort, L. Olender, O. Beaven, X. Huang, E. J. Brown, M. Vanden Bempt, H. M. Khoo, J. Bhadury, T. A. Milne, A. C. Wilkinson, A genome-wide screen identifies Runx2 as a novel regulator of hematopoietic stem cell expansion and T-cell commitment. Blood 146, 3188–3200 (2025).

81. J. Liu, D. K. Kumar, B. Hurieva, F. Jonas, N. Barkai, Engineering intrinsically disordered regions for guiding genome navigation. Mol. Cell 86, 851–867.e5 (2026).

82. J. Yan, M. Enge, T. Whitington, K. Dave, J. Liu, I. Sur, B. Schmierer, A. Jolma, T. Kivioja, M. Taipale, J. Taipale, Transcription factor binding in human cells occurs in dense clusters formed around cohesin anchor sites. Cell 154, 801–813 (2013).

83. Cell Editorial Team, Moving structural biology forward together. *Cell* 187, 511–512 (2024).

84. 84. A. Shanehsazzadeh, Autonomous De Novo Protein Binder Design with Claude. *alphaXiv* (2026).

85. J. B. Grimm, L. Xie, J. C. Casler, R. Patel, A. N. Tkachuk, N. Falco, H. Choi, J. Lippincott-Schwartz, T. A. Brown, B. S. Glick, Z. Liu, L. D. Lavis, A general method to improve fluorophores using deuterated auxochromes. JACS Au 1, 690–696 (2021).

86. J. B. Grimm, B. P. English, H. Choi, A. K. Muthusamy, B. P. Mehl, P. Dong, T. A. Brown, J. Lippincott-Schwartz, Z. Liu, T. Lionnet, L. D. Lavis, Bright photoactivatable fluorophores for single-molecule imaging. Nat. Methods 13, 985–988 (2016).

87. M. Tokunaga, N. Imamoto, K. Sakata-Sogawa, Highly inclined thin illumination enables clear single-molecule imaging in cells. Nat. Methods 5, 159–161 (2008).

88. A. D. Edelstein, M. A. Tsuchida, N. Amodaj, H. Pinkard, R. D. Vale, N. Stuurman, Advanced methods of microscope control using Î¼Manager software. J. Biol. Methods 1, e10 (2014).

89. D. B. Allan, T. Caswell, N. C. Keim, C. M. van der Wel, R. W. Verweij, Soft-Matter/Trackpy: Trackpy v0.5.0 (Zenodo, 2021; 10.5281/zenodo.4682814).

90. A. Heckert, L. Dahal, R. Tjian, X. Darzacq, Recovering mixtures of fast-diffusing states from short single-particle trajectories. Elife 11, e70169 (2022).

91. M. Bulajić, D. Srivastava, J. S. Dasen, H. Wichterle, S. Mahony, E. O. Mazzoni, Differential abilities to engage inaccessible chromatin diversify vertebrate Hox binding patterns. Development 147, dev194761 (2020).

92. S. G. Landt, G. K. Marinov, A. Kundaje, P. Kheradpour, F. Pauli, S. Batzoglou, B. E. Bernstein, P. Bickel, J. B. Brown, P. Cayting, Y. Chen, G. DeSalvo, C. Epstein, K. I. Fisher-Aylor, G. Euskirchen, M. Gerstein, J. Gertz, A. J. Hartemink, M. M. Hoffman, V. R. Iyer, Y. L. Jung, S. Karmakar, M. Kellis, P. V. Kharchenko, Q. Li, T. Liu, X. S. Liu, L. Ma, A. Milosavljevic, R. M. Myers, P. J. Park, M. J. Pazin, M. D. Perry, D. Raha, T. E. Reddy, J. Rozowsky, N. Shoresh, A. Sidow, M. Slattery, J. A. Stamatoyannopoulos, M. Y. Tolstorukov, K. P. White, S. Xi, P. J. Farnham, J. D. Lieb, B. J. Wold, M. Snyder, ChIP-seq guidelines and practices of the ENCODE and modENCODE consortia. Genome Res. 22, 1813–1831 (2012).

93. H. Li, Aligning sequence reads, clone sequences and assembly contigs with BWA-MEM, arXiv [q-bio.GN] (2013). 10.48550/arXiv.1303.3997.

94. H. Li, B. Handsaker, A. Wysoker, T. Fennell, J. Ruan, N. Homer, G. Marth, G. Abecasis, R. Durbin, 1000 Genome Project Data Processing Subgroup, The Sequence Alignment/Map format and SAMtools. Bioinformatics 25, 2078–2079 (2009).

95. A. R. Quinlan, I. M. Hall, BEDTools: a flexible suite of utilities for comparing genomic features. Bioinformatics 26, 841–842 (2010).

96. F. Ramírez, D. P. Ryan, B. Grüning, V. Bhardwaj, F. Kilpert, A. S. Richter, S. Heyne, F. Dündar, T. Manke, deepTools2: a next generation web server for deep-sequencing data analysis. Nucleic Acids Res. 44, W160–5 (2016).

97. P. Ewels, M. Magnusson, S. Lundin, M. Käller, MultiQC: summarize analysis results for multiple tools and samples in a single report. Bioinformatics 32, 3047–3048 (2016).

98. J. Feng, T. Liu, B. Qin, Y. Zhang, X. S. Liu, Identifying ChIP-seq enrichment using MACS. Nat. Protoc. 7, 1728–1740 (2012).

99. Q. Li, J. B. Brown, H. Huang, P. J. Bickel, Measuring reproducibility of high-throughput experiments. Ann. Appl. Stat. 5, 1752–1779 (2011).

100. R. Stark, G. Brown, *DiffBind: Differential Binding Analysis of ChIPSeq Peak Data* (Cancer Research UK -Cambridge Institute, Cambridge, 2026).

101. M. I. Love, W. Huber, S. Anders, Moderated estimation of fold change and dispersion for RNA-seq data with DESeq2. Genome Biol. 15, 550 (2014).

102. F. Pedregosa, G. Varoquaux, A. Gramfort, V. Michel, B. Thirion, O. Grisel, M. Blondel, A. Müller, J. Nothman, G. Louppe, P. Prettenhofer, R. Weiss, V. Dubourg, J. Vanderplas, A. Passos, D. Cournapeau, M. Brucher, M. Perrot, É. Duchesnay, Scikit-learn: Machine Learning in Python, arXiv [cs.LG] (2012). 10.48550/arXiv.1201.0490.

103. B. Langmead, C. Trapnell, M. Pop, S. L. Salzberg, Ultrafast and memory-efficient alignment of short DNA sequences to the human genome. Genome Biol. 10, R25 (2009).

104. A. Dobin, C. A. Davis, F. Schlesinger, J. Drenkow, C. Zaleski, S. Jha, P. Batut, M. Chaisson, T. R. Gingeras, STAR: ultrafast universal RNA-seq aligner. Bioinformatics 29, 15–21 (2013).

105. Y. Liao, G. K. Smyth, W. Shi, featureCounts: an efficient general purpose program for assigning sequence reads to genomic features. Bioinformatics 30, 923–930 (2014).

106. Y. Benjamini, Y. Hochberg, Controlling the false discovery rate: A practical and powerful approach to multiple testing. J. R. Stat. Soc. Series B Stat. Methodol. 57, 289–300 (1995).

107. A. P. Thompson, H. M. Aktulga, R. Berger, D. S. Bolintineanu, W. M. Brown, P. S. Crozier, P. J. in ’t Veld, A. Kohlmeyer, S. G. Moore, T. D. Nguyen, R. Shan, M. J. Stevens, J. Tranchida, C. Trott, S. J. Plimpton, LAMMPS -a flexible simulation tool for particle-based materials modeling at the atomic, meso, and continuum scales. Comput. Phys. Commun. 271, 108171 (2022).

108. R. Gowers, M. Linke, J. Barnoud, T. Reddy, M. Melo, S. Seyler, J. Domański, D. Dotson, S. Buchoux, I. Kenney, O. Beckstein, “MDAnalysis: A python package for the rapid analysis of molecular dynamics simulations” in Proceedings of the Python in Science Conference (Los Alamos National Laboratory (LANL), 2016), pp. 98–105.

109. N. Michaud-Agrawal, E. J. Denning, T. B. Woolf, O. Beckstein, MDAnalysis: a toolkit for the analysis of molecular dynamics simulations. J. Comput. Chem. 32, 2319–2327 (2011).

110. R. J. Emenecker, D. Griffith, A. S. Holehouse, Metapredict: a fast, accurate, and easy-to-use predictor of consensus disorder and structure. Biophys. J. 120, 4312–4319 (2021).

111. N. Morffy, L. Van den Broeck, C. Miller, R. J. Emenecker, J. A. Bryant Jr, T. M. Lee, K. Sageman-Furnas, E. G. Wilkinson, S. Pathak, S. R. Kotha, A. Lam, S. Mahatma, V. Pande, A. Waoo, R. C. Wright, A. S. Holehouse, M. V. Staller, R. Sozzani, L. C. Strader, Identification of plant transcriptional activation domains. Nature 632, 166–173 (2024).

112. M. Ashburner, C. A. Ball, J. A. Blake, D. Botstein, H. Butler, J. M. Cherry, A. P. Davis, K. Dolinski, S. S. Dwight, J. T. Eppig, M. A. Harris, D. P. Hill, L. Issel-Tarver, A. Kasarskis, S. Lewis, J. C. Matese, J. E. Richardson, M. Ringwald, G. M. Rubin, G. Sherlock, Gene ontology: tool for the unification of biology. The Gene Ontology Consortium. Nat. Genet. 25, 25–29 (2000).

113. N. Alerasool, H. Leng, Z.-Y. Lin, A.-C. Gingras, M. Taipale, Identification and functional characterization of transcriptional activators in human cells. Mol. Cell 82, 677–695.e7 (2022).

